# Modelling non-local neural information processing in the brain

**DOI:** 10.1101/2022.01.27.477993

**Authors:** Johannes Balkenhol, Barbara Händel, Juan Prada, Conrado A. Bosman, Hannelore Ehrenreich, Johannes Grohmann, Jóakim v. Kistowski, Sonja M. Wojcik, Samuel Kounev, Robert Blum, Thomas Dandekar

## Abstract

The representation of the surrounding world emerges through integration of sensory information and actions. We present a novel neural model which implements non-local, parallel information processing on a neocolumnar architecture with lateral interconnections. Information is integrated into a holographic wave interference pattern. We compare the simulated *in silico* pattern with observed *in vivo* invasive and non-invasive electrophysiological data in human and non-human primates. Our model replicates the modulation of neural high-frequency activity during visual perception showing that phase-locked low and high-frequency oscillations self-organize efficiently and carry high information content. The simulation further models how criticality (high content) of information processing emerges given a sufficiently high number of correlated neurons. Non-local information processing, forming one holographic wave pattern, suggests a platform for emergence of conscious perception.

**One sentence summary:** Simulated non-local information processing on a neocolumnar architecture models well multiple electrophysiological observations of brain activity, including high-frequency activity during visual perception in primates.

## Introduction

The human brain relies on the interplay of neuronal circuits to form a network underlying consciousness, defined as subjective experience. Such interplay has been shown to include serial processing and neuronal recognition as well as integrative properties and holistic processes ^1, 2^. Further, coordinated firing and synchronous synaptic activity of neurons are typical elements of higher order neuronal mechanisms ^3^, representing information processing in the brain that correlates with experience ^4, 5^. Low and high frequency phenomena at the single cell level up to neural networks, including oscillatory patterns in postsynaptic potentials and firing activity, can contribute to cellular or synaptic plasticity and thereby shape learning and memory ^3, 6^ and many other cognitive processes such as perception ^7^.

Processing of sensory input in neurons and neuronal circuits have been well defined over the last decades (Buzsaki et al., 2013; Odegaard et al., 2017; Tononi et al., 2016). Considerably less is known about how different sensory input is integrated and combined with the current physical (motor) state and the current cognitive state, including memory or integrated perception and, ultimately, consciousness. To achieve an integration of sensory and motor processes, to explain actions and probe consciousness, models have been developed that capture information generated in neuronal circuits. Such models elucidate the unique integrative properties of conscious perception, by integrating sensory elements as well as voluntary actions ^1, 5, 8^.

Here we asked whether non-local information processing may be at the root of cortical information processing. We therefore built up a non-local processing model based on a neocolumnar architecture ^2, 9^ with lateral connections between the columns. Here, complexity is generated by repeating simple rules in time and space, which pins down the underlying process of emergence. Exploiting the advances in computing power for large-scale grid computing allowed us to apply these simple rules to large networks. The simulation shows how high frequency pattern encode high information content. This high frequency coded information, modulated in a wave-like fashion through the lateral connections in the neocolumns, reaches all participating columns, thereby creating a holistic representation. A high enough number of neurons, however, turns out to be essential to form stable high information content. Importantly, we can demonstrate the phase-locked high and low frequency (HF and LF) pattern implied by the simulation in multielectrode and ECoG (electrocorticography) brain recordings in monkeys performing a visual perception task.

## Results

In biological systems, feedback loops exist on the level of DNA, signaling cascades, cellular or neuronal networks (**fig. S1 and fig. S2**). In cells, genes encode proteins; these proteins regulate cellular phenotypes in signaling cascades that are determined by an interplay of positive and negative feedback (**fig. S2**). For computational modelling of cells, this information is used to describe how cellular phenotypes emerge from an interplay of modular signaling compounds (signaling pathways, as exemplarily shown in **fig. S3**).

We asked whether the reduction of the architecture of a cellular simulation on a unified model of activation and inhibition could create a new emergent level of information processing. For this, we created a simulation based on laterally interconnected microcircuits, a model for non-local information processing.

The kernel of the model is shown in **eq. 1**. Computational processing steps of the simulation are outlined in **Fig. 1A** (see extended description in Material and methods).

**Figure 1:**
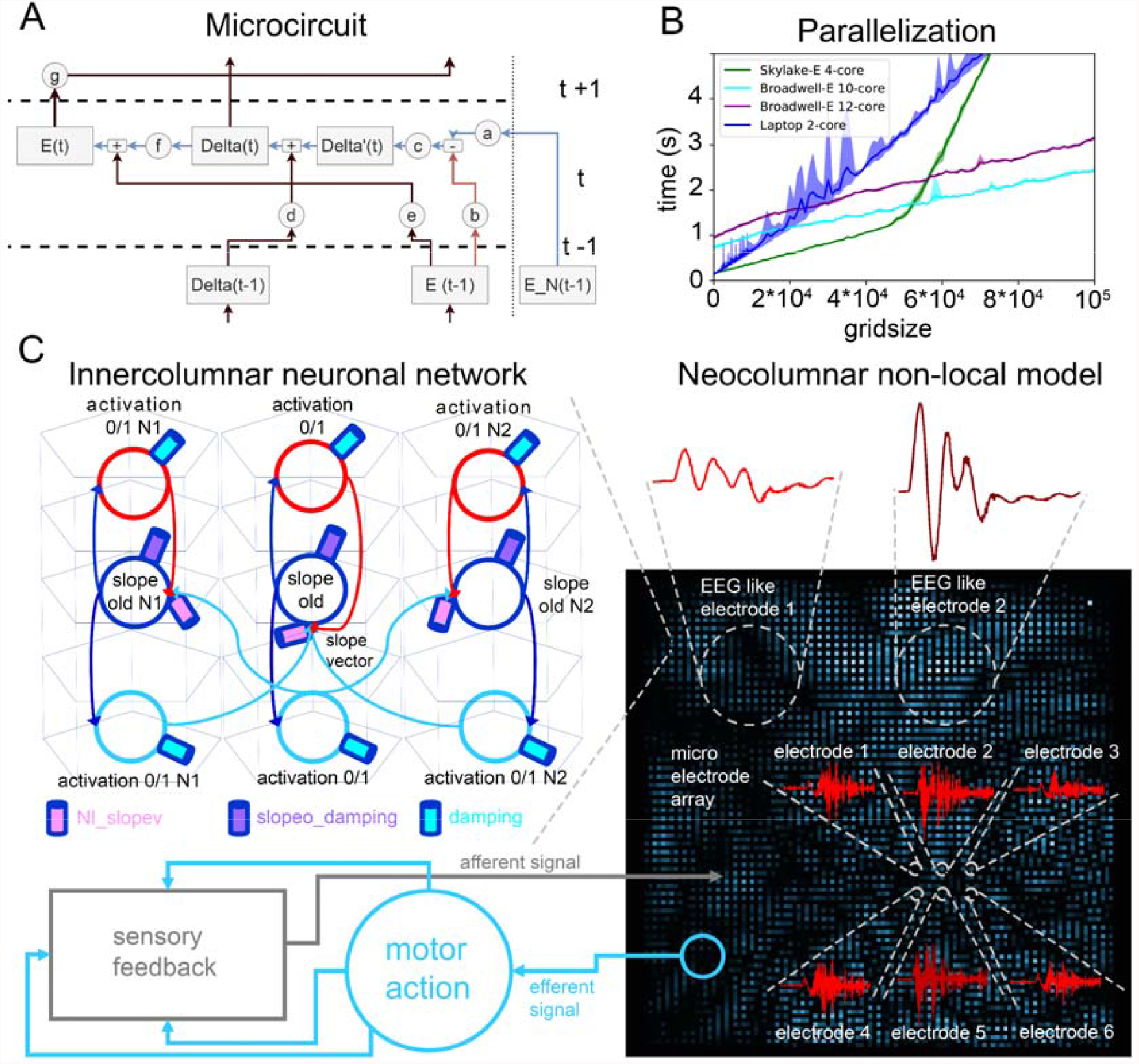
A non-local information processing model is embedded in a neocolumnar architecture. **(A)** The non-local information processing model relies on the unification of an interplay of activation and inhibition in a microcircuit and their interconnections. **(B)** A computer cluster simulation of non-local processing. Shown is a simulation of non-local information processing and its dependence on the number of simulated microcircuits in a grid (gridsize). The information was processed more efficiently with more nodes but there were no emergent new properties. **(C)** The neocolumnar non-local information processing model combines neocolumns as microcircuits and interconnects them laterally. The organization of microcircuits in a grid of neocolumnar topology is the basis of the neocolumnar non-local information processing model. On this processing platform, the information was represented as a whole, and complex wave interference pattern arise. The information, such as motor action and feedback from the world, is distributed over the entire model, thus available at any column over time. The information is encoded in frequency and phase. Phenomena like fast ripple-like energy changes are formed. ERP-like signals appear as sum of multiple ripple-like signals. Here, each pixel represents one neocolumnar circuit, which is also valid for the recorded ripple-like activity. The dashed lines indicate size relations.

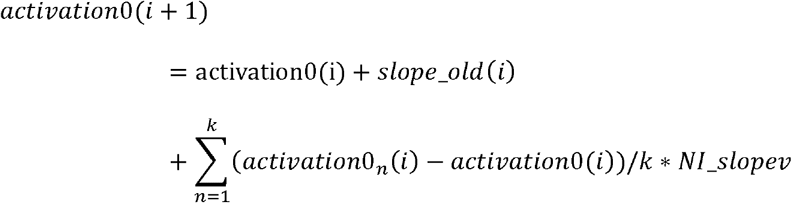

When we computed the simulation, input signals were convoluted, copied, and spread over the entire model space (**fig. S4**). This enabled the interference of all incoming signals providing “the whole of the information”, as postulated for holography ^10^. This created a new level of emergence. Oscillating wave-like signals self-organized and appeared in face of simple information input to the simulation (**fig. S4**).

To probe any new higher emergence level, a high number of our simulations were coupled on a large computer cluster. Parallel computing sped up the simulations, exhibiting constant performance due to the linear increase of computation time based on the number of nodes (**Fig. 1B**). We simulated up to 400,000 nodes in real time using a resolution of one tick per millisecond on a 24-core server system. Processing time increased linearly with the grid size. However, further processing power did not lead to new emergence levels but led only to linear gains and losses of processing power versus communication overhead (**Fig. 1B**).

The analysis of the wave-like signals (**fig. S5**) showed similarities to the critical distribution as defined by the Ising model ^11^ and as found in recordings of brain activity ^12-14^. Furthermore, is has been suggested that critical distributions indeed represent a state of maximal integrated information processing ^15^. In our simulation, criticality was stabilized by the model architecture and respective energy coupling parameters (**eq. 1**) and was robust over a wide range of energy coupling modulations (**fig. S5**). This robustness is in contrast to fragile critical states in the Ising model, which describes the self-organization of complex second-order phase transitions between the homogeneous states of order (subcritical) and chaos (supercritical; **fig. S5**, see also ^13^).

We found that many properties of our model were in accordance with observations in the cortex. For instance, microcircuits are typical units of information processing in the brain ^3^ and critical distributions have also been found in the mammalian cortex ^12-14^. Therefore, we asked whether a simulated neocolumnar architecture ^2, 9^ would also allow wave-like non-local information processing.

For generating an innercolumnar network with lateral connections, we used a simple neuronal oscillator design (**Fig. 1C**), consisting of two types of neurons (inhibitory and excitatory) and allowing summation as well as subtraction. In the neocolumnar analogue, we combined excitatory and inhibitory neurons in a feedback loop (representative for the innercolumnar network) modulated by the neighboring columns (**Fig. 1C**, the interconnection). The interconnections build a spatial derivative of the neighboring energy levels (*activation1N1-4*) to the previous energy level of the column in the center (*activation0/1* in red). The spatial derivation is followed by two integration steps of excitatory neurons (*slope_old* and *activation0/1* in blue) in order to represent temporal integration (**Fig. 1C**). In sum, the addition, and the subtraction of **eq. 1** is here transformed into an interplay of excitatory and inhibitory neurons. The modulation of the energy transfer between the neurons, such as electrical or chemical communication within the microcircuit analogue, is incorporated by the energy coupling parameter *NI_slopev, slopeo_damping* and *damping* (**Fig. 1C**). Here, *slopeo_damping* and *damping* represent neural mechanisms that can dampen the persistent energy level of neurons, as it happens, for instance, at the cell body of neurons by the action of the neurotransmitter GABA. The central energy coupling parameter, *NI_slopev*, modulates the energy transmission of the transient energy state of a center node versus its neighboring nodes (spatial derivation; see **eq. 1**). A decrease *NI_slopev* represents a damping of the energy transmission, whereas a decrease of *slopeo_damping* and *damping* facilitate energy transmission. In the simulation of the neuronal microcircuit and in the neocolumnar neuronal network, *NI_slopev* represents the simulated transmission of energy between neurons via synapses, meaning the efficacy of excitatory and inhibitory synaptic transmission.

After constructing the neocolumnar neuronal network (**Fig. 1C**), we transferred parameters, such as diameter, processing speed, timing, and topology from the neocolumnar architecture to the neocolumnar non-local information processing simulation combining 14,400 neocortical columns (see Materials and methods). If a simple, rhythmic peak signal was processed in the simulation, the input was time-dependently transformed into a holographic wave interference pattern that provided all individual neocortical columns with the same frequency information (see **suppl. video S3** and **fig. S4**).

Other input signals can also be coded into energy level modulations, over time and space. By design, sensory feedback from the environment is provoked by efferent signals, e.g. motor action (in blue in **Fig. 1C**). The sensory feedback is represented by afferent sensory signals (in grey in **Fig. 1C**) that interfere with the existing information in the model (**Fig. 1C**).

To investigate how different signal input behaves in the model, we considered chaotic, periodic-peak, sinusoid, and complex signals (**suppl. video S4**). To simulate broad and active processing in the brain, we used a complex stimulus consisting of short bursts (50 – 100 ms) of high frequency (300 – 500 Hz) superimposed on a reference wave with lower frequency (1 - 20 Hz) as a reference input signal. When we applied more than 50 of these complex stimuli, with a random onset, a complex interference pattern and a holistic representation of the stimuli in time and space appeared (**Fig. 1C**) (see **suppl. video S5**).

For the read out, the information processed in the model can be extracted at every point in phase and frequency and can again be offered to the model (interface design). As any kind of input information for the simulation is processed into frequency and phase (**Fig. 1C**), we used a Fourier transform for the decoding and read out of the simulated signals. To extract the processed information content from the simulation, the model uses virtual electrodes of different size (as exemplarily indicated by white circles in **Fig. 1C**). The user can define the electrode diameter in number of processing units (columns). Extraction of the information on its smallest scale is given by the diameter of one column ^2, 9^. In accordance with Mountcastle (1997), the diameter of the virtual electrode was set to 500 μm to represent one neocortical column. Notably, extraction of the signal induced by the complex stimulus described above (reference signal), with such a small virtual electrode resulted in signals that are primarily composed of fast ripple-like energy changes (**Fig. 1C**, shown by electrode 1 - 6). The dominating fraction of fast oscillations in small electrode signals is similar to those described from recordings using microelectrode arrays ^16^. By increasing the diameter of the virtual electrode, thus measuring multiple columns and ripple-like events at the same time, the extracted simulated signal looked like typical event-related brain potentials (ERP) (**Fig. 1C**).

We used virtual electrodes of different diameter (500 μm, 2 mm, 2 cm) to simulate different electrophysiological recordings (microelectrode recording, as used in MEA; field electrode as used in EEG) and signals integrated over multiple neocolumns (representative for LFP, ERP-like signals). These simulations show that our model can mimic a broad range of electrophysiological signals and energy phenomena such as critical distributions, harmonics (overtones), coherence patterns, self-organized signal oscillations, and stimulus-induced high-frequency oscillations (**suppl. table 2**).

Further, we tested how the model behaves when the energy transmission, representing changes in synaptic communication, was modulated. A *NI_slopev* of 0 defines the lower border for information processing and the input signal remains unmodulated. Above the critical *NI_slopev* value of 2.6655 the model collapses. In the range of *NI_slopev* between > 0 and 2.6655, signals can be processed in frequency and phase in this model, with its given size.

We tested the model with complex input (see **Fig. 1C**) and simulated EEG-like read outs. With the parameter *NI_slopev* (**eq. 1**) at 2.6655, representing the upper limit of the model, the output signal shows fast oscillations that are small in amplitude (**Fig. 2A**). Shifting the value to 0.1, slow oscillatory signals with larger amplitudes were formed. A decrease of inhibitory components in the model, given by the values *slopeo_damping* (1.0E-4→1.0E-5) and *damping* (1.0E-2→1.0E-3), slightly modulated the amplitudes of the virtual waves and facilitated the formation of regular slow waves. We define these states as waking model state (upper border of the model; **suppl. video S6**) or slow wave sleep model state (SWS; **suppl. video S7**) (lower border of the model). Based on these definitions (**Fig. 2A**), we compared simulated model states with published data (**suppl. table 2**). First, we tested the peak distributions of virtual MEA signals of the waking and the SWS state. As shown in **Fig. 2B**, both distributions could be fitted with a lognormal function, as seen earlier in biological MEA recordings of waking and SWS states of the rat brain ^14^. Starting from the modelled waking state, an anesthesia-like state (**Fig. 2C** and **suppl. video S8**) could be computed by an increase of the inhibitory influence to virtual cell bodies (inhibitory parameters: *slopeo_damping* (1.0E-2 → 1.0E-1) and *damping* (1.0E-4 → 1.0E-3)). Here, synaptic transmission parameter *NI_slopev* was kept constant. We could also observe the transformation from a lognormal peak distribution to a power law distribution at the onset of the anesthesia-like state (**Fig. 2C)** as experimentally observed in ^14^).

**Figure 2:**
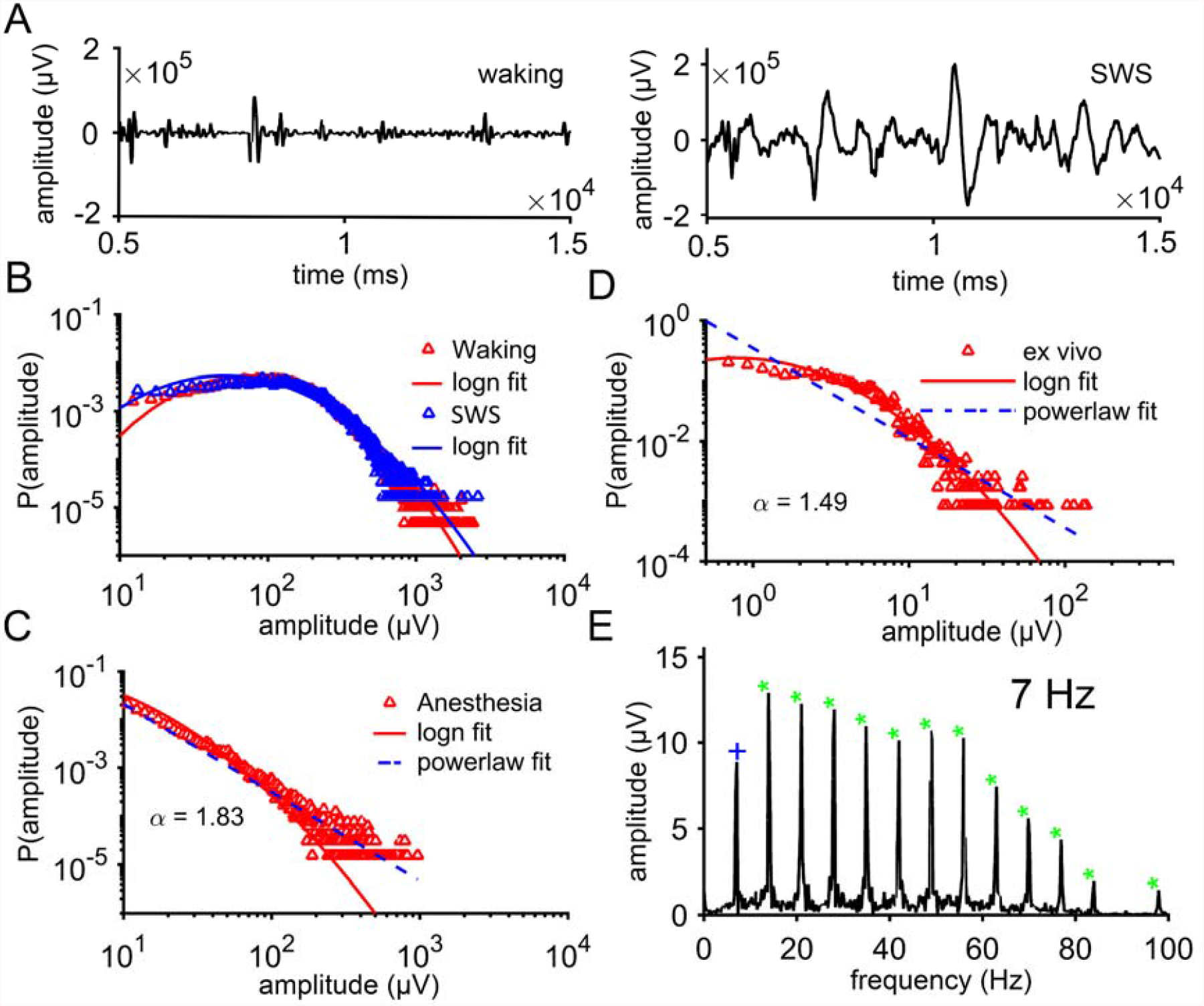
Analysis of the neocolumnar non-local information processing simulation. Distinct characteristics of the simulations are rooted by the energy transmission of the innercolumnar and lateral connections. Fundamental energy features are also found in electrophysiological recordings of the brain and can be utilized to discriminate brain states. **(A)** A complex input to the model reproduces EEG-like signals representative for a waking state. The decrease of the energy coupling parameter *NI_slopev* only, could switch the model into the simulated slow wave sleep (SWS) state. **(B)** The peak distribution of MEA signals at waking and SWS were fitted by a lognormal distribution and thus could indicate criticality. **(C)** When simulating anesthesia by increasing the *damping* and *slopeo_damping* simulating damping of the cell bodies the lognormal distribution morphed into a power law distribution. The shift from a lognormal to a power law fit indicates an increased localization of the signal processing. **(D)** Similar lognormal distributions were also found in electrode recordings of hippocampal brains slice of mice. Here, the power law fit did not apply. **(E)** Harmonics self-organize in the model in response to periodic rectangular input and were visualized in a simulated LFP (shown), EEG and MEA recordings. The root frequency is shown by a blue cross and the resulting harmonics are indicated by green stars.

Subsequently, we asked whether critical distribution values, as indicated by our modelling, would also appear in *ex vivo* electrode recordings from hippocampal slices of mice. For the hippocampus, the peak distribution in signals from field recordings could also be fitted with a lognormal function (**Fig. 2D** and **fig. S6**). This shows a similarity between modelled signal states of waking and SWS and corresponding field recording signals from hippocampal slices (**Fig. 2D**). The anesthesia-like model state, indicated by a power law, was not found in the biological data (**Fig. 2D**). Notably, we observed the emergence of harmonics in response to periodic peak signals in the model (**Fig. 2E**). These harmonics decline in the simulation when the signal is sinusoid (**fig. S7**). Both model findings have been observed in *in vivo* MEA recordings ^17^.

Next, we examined the parameters of our simulation more closely, matching experimental data are referenced and summarized in the supplement (in particular Table S2). First, we asked, how many frequencies we can encode in the waking state, with a maximum value of synaptic transmission (*NI_slopev)*. As shown in **Fig. 3A**, it was possible to decode more than 150 different frequencies and corresponding harmonics, within 3 s, with one virtual electrode, from one simulated neocortical column. This temporal aspect in signal emergence corresponds nicely to consciously processed visual stimuli (50 bits per second) integrated over 3 s ^18^. In addition, the half-width of the peaks (in Hz) allows to estimate the coding potential of the simulation. As the half-width was determined to be ∼0.5 Hz within 3 seconds (see **fig. S8**) the model could code up to ∼329 bit/s in a bandwidth of 7 - 500 Hz at a single location. Moreover, we showed that the value of synaptic transmission (*NI_slopev*) defines the maximum frequency that can be processed and thereby shows a linear correlation to the speed of the travelling waves (**fig. S9**), as suggested by observed data earlier ^19^. In turn, fast travelling waves are a clear signature of high information processing.

**Figure 3:**
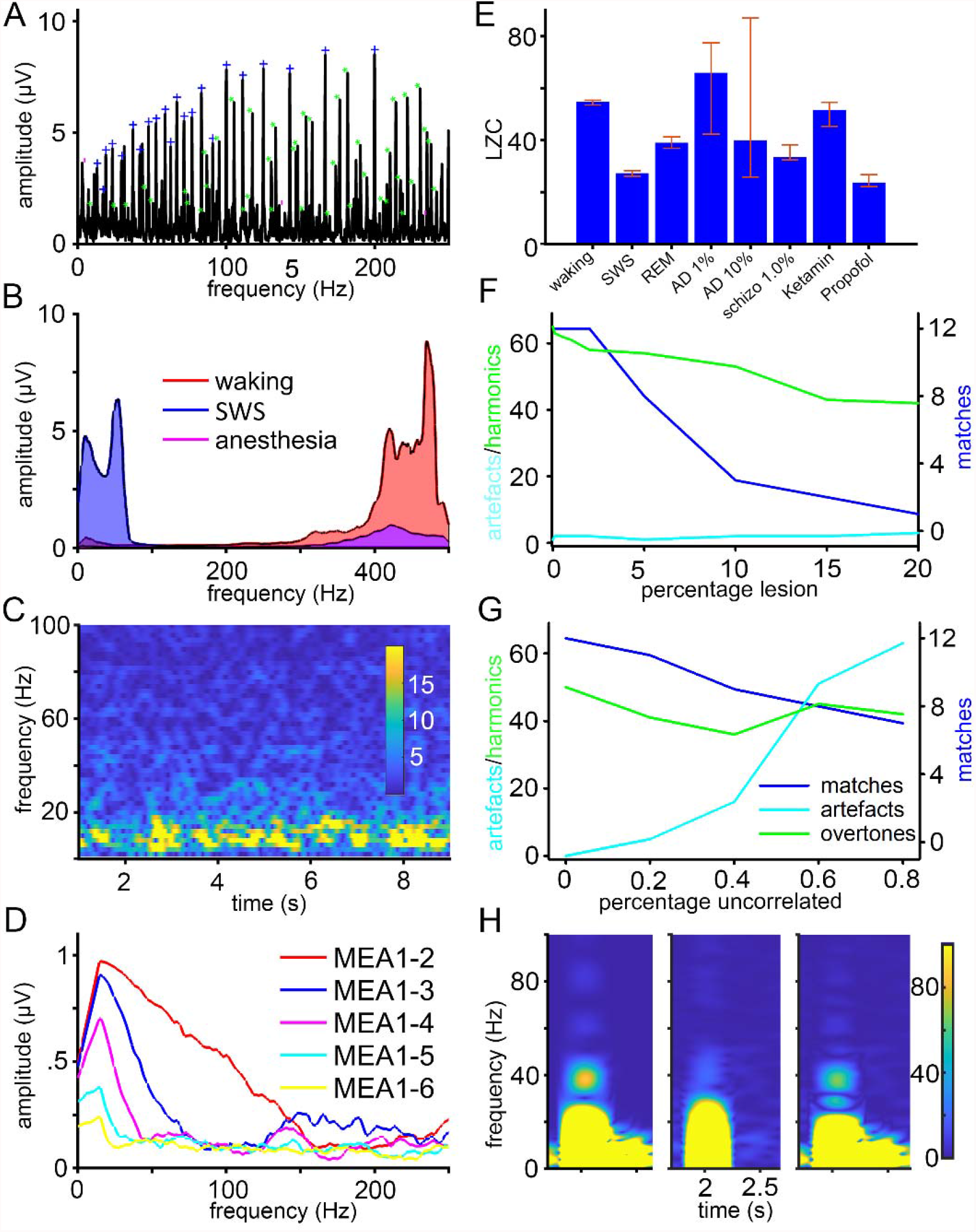
*In silico* simulation parameter effects on neocolumnar non-local processing of stimuli. **(A)** Efficient resolution of more than 50 stimuli as complex model input indicates the prove of concept of processing frequency information (blue cross: root frequency; green: harmonics). **(B)** The resonating frequency band of MEA signals during simulated waking, SWS and anesthesia. We show different favored bandwidth according to the respective activation level. **(C)** Self-organizing baseline activity of the model at spontaneous activation mimics observations of EEG baselines. **(D)** Coherence decreases with frequency and distance when spontaneous activity is the only input. **(E)** Lempel Ziv complexity of simulated EEG recordings. The analysis visualizes complexity of information processing and is used to discriminate different simulated brain states. LZC values are stated as median of six electrodes and error bars indicate the 15% and 85% quantile. **(F)** Loss of neocolumns is used to mimic information processing in Alzheimer’s disease-like state. Processing of information remains robust, however, with an increasing loss of neocolumns, the spread of information processing is impaired. **(G)** In simulated schizophrenia, a low correlation of energy transmission between neurons decreases frequencies that match to the input frequencies (matches). Frequencies that did not match to the input frequencies or the respective harmonics increased (artefacts). **(H)** In modelled schizophrenia, a gamma-band decline is observed. The self-organizing gamma-band following a stimulus is superimposed on a reference and is shown for a simulated healthy (left) neocortical model and simulated schizophrenia (middle). (right) is the delta of both models. The color bar indicates simulated activity in μV.

The control of synaptic transmission (*NI_slopev)* on maximal coding becomes more evident when outlining the resonating frequencies bands of simulated MEA signals of the states waking, slow wave sleep and anesthesia, at least when chaotic input is applied. The waking state aligns more in the high frequency (HF) bandwidth and the SWS aligns more in the low frequencies (LF) bandwidth.

In effect, the decrease of the synaptic transmission (*NI_slopev* function) acts as a low pass filter and reduces HF coding. Finally, anesthesia seems to suppress HF and LF coding likewise (**Fig. 3B)**.

Within these ranges, model size has no influence on the direct decoding of the input frequencies but affects the self-organization of harmonics (**fig. S10**) allowing increased number of stable frequencies with increased model size. This becomes obvious when analyzing the resonating frequencies at different model sizes for spontaneous activity only. Resonating frequencies and thereby the number of eigenstates drastically increases with model size (**fig. S11**). Importantly, the increase in model size potentially enhances phase coding and this enables more frequencies to be coded in parallel (**fig. S11**).

Following, the self-organizing effect of the model, simulated, spontaneous activity of cortical areas organizes after several seconds without external stimuli at around ∼8 Hz at waking (**Fig. 3C**) and decreases to theta activity in the model SWS state (**fig. S12**). This matches well will other experimental observations of baseline activity or recordings during SWS (see **table S2**). Furthermore, we found that coherence between simulated MEA electrodes changes during spontaneous baseline activity dependent on frequency and distance (**Fig. 3D**). The gradual decline of coherence with increasing frequency and distance between the electrodes matches with cortical measurements of visual perception tasks (see **table S2**). However, sinusoid stimuli can increase the coherence also at higher distances and frequencies, this counteracts the coherence decline in simulation (**fig. S13**) and experiment (see **table S2**).

We created other model states by fine-tuning of model parameters. Specifically, we faithfully reproduced anesthesia (Ketamin, Propofol; **suppl. video S8** and **S9**), a rapid-eye-movement sleep state (REM; **suppl. video S10**) and disease states including Alzheimer’s disease (AD; **suppl. video S11** and **S12**) and schizophrenia (**suppl. video S13**). We compared the complexity of the resulting simulated wave patterns using the Lempel Ziv compression (LZC; **Fig. 3E)**. The complexity measure helped to differentiate between high complex signal processing and low complex signal processing. Our comparison of states suggests increased information processing during the waking state or information decline for the SWS, anesthesia, and REM states. In AD, a high variation of LZC values indicates local differences in complexity compared to healthy states. In schizophrenia, the LZC value was dependent on the percentage of uncorrelated energy coupling between simulated neocortical columns. Overall, LZC values compared well to experimentally observed measurements (**table S2**).

The pathology of AD ^20^ show a robustness against lesions or defects. To simulate lesions, we randomly inactivated neocolumns from the model. The simulation could still decode full information when 4% of all neocolumns were lesioned. Even when 20% of the neocolumns were lesioned, information decoding was still present (**Fig. 3F**), however, it appeared rather localized (**suppl. video S11** and **S12**). Schizophrenia was simulated by lowering the correlation of synaptic energy transfer between the neurons.

The energy transmission (parameters *NI_slopev, slopeo_damping* and *damping*) of some of the neighboring columns was randomly impaired causing unsymmetrical processing. We could show that the model is sensitive to uncorrelated processing as this causes the generation of artificial frequencies that do not correlate with the input frequencies (**Fig. 3G**). In our simulation (**Fig. 3H**) we could replicate uncorrelated neuronal signaling and a distinct pathology phenomenon of schizophrenia, the decline of the evoked gamma-band ^21, 22^. Following a LF-coupled HF stimulus, the recorded processed signal was composed of a self-organizing theta and gamma-band (**Fig. 3H, left)**. In the schizophrenia model (**Fig. 3H, middle)**, we see a decline of the evoked gamma-band in response to a complex stimulus (**Fig. 3H, right)** in line with observations ^21-23^. Especially, uncorrelated damping (parameter *slopeo)* decreases the emergence of self-organized gamma-band in the model.

Other phenomena modelled in our simulation, such as beta band firing (**fig. S14**), simulation of epilepsy (**fig. S15**) and the effect of spatial under-sampling in large electrodes masking HF signaling (**fig. S16**) fit again well with observations and are discussed in the supplemental material.

*In silico*, the simulation provides evidence for HF information coding and LF coupling that self-organizes due to specific resonance properties. If high frequencies are indeed at the basis of *in vivo* information transmission, this would suggest HF signals in close to all electrophysiological recordings. Accordingly, we analyzed the neural response to visual input (see the used grating stimulus in **fig. S17B)** in *in vivo* macaque V1 recordings and compared it to our *in silico* model output (**fig. S17A** and **suppl. video S14)** in response to comparable input ^24, 25^.

Here it is important to consider the different *in vivo* recording methods. Microelectrodes pick up neural activity from responding neurons at the site of stimulation, as secured by receptive field (RF) mapping. Therefore, these neural responses mainly depict incoming sensory information, which would be, according to our simulation (**fig. S18**), encoded in the HF signal and distributed via slower waves to neighboring sites. ECoG, which applies larger electrodes placed subdurally, picks up neural activity from a larger area. If placed over the site of stimulation, the recording includes signal from responding neurons as well as surrounding neurons, not directly driven by the input. As we assume a lateral distribution of information via low frequency waves, a HF/LF phase relationship is predicted to be particularly visible in those recordings. To test these considerations by experiment, we compared V1 multielectrode recordings from neurons that were driven by the sensory input to model data from the site of stimulation. Furthermore, we compared ECoG recordings over V1 with model data from an area surpassing the site of stimulation. For both, biological and model data, we could observe the predicted pattern of HF signals coupled to LF signals, self-organized in face of frequency unspecific stimulation (**Fig. 4**). Specifically, the V1 multielectrode recordings (RFs overlapping with the stimulus) showed a broadband HF (200-1000 Hz) power increase after stimulus onset, comparable to the simulated data generated for the site of sensory input (**Fig. 4A**). As could be shown in the animal data, this induced HF increase largely corresponded to the observed modulation of multiunit and spiking activity, respectively (**Fig. 4C**). The evoked activity is defined as averaged activity over stimulus onsets thereby highlighting the time-locked modulation. Importantly, the evoked HF activity showed a temporal pattern independent from MUA and LFP (**Fig. 4B**). Only this evoked activity was modulated in its power by a slow phase (**Fig 4D**). If we assume that this time locked HF modulation depicts the wave-like lateral distribution of information, we should find this slow power modulation particularly in neurons surrounding the ones processing the stimulus. Analyzing ECoG recordings from subdural electrodes placed over V1, picking up neural activity from the site of visual processing but additionally from a surrounding area, we indeed find HF increase for induced and evoked data (**Fig. 5A and B**). Importantly, the induced HF pattern was already phase modulated similar to the evoked pattern (**Fig. 5C and D**). Model data generated for sites only partly overlapping with the sensory input was again comparable between the simulation and biological data (**Fig. 5**). The topography of the stimulus-evoked and stimulus-induced HF power changes further showed that induced HF power modulation was confined to V1 (**Fig. S17C**). The evoked HF power change spread to V2, supporting the idea of information transfer in a temporally correlated fashion (**Fig. S17D**). We gave a preliminary report of HF modulation in the above described electrophysiological recordings ^26^.

**Figure 4:**
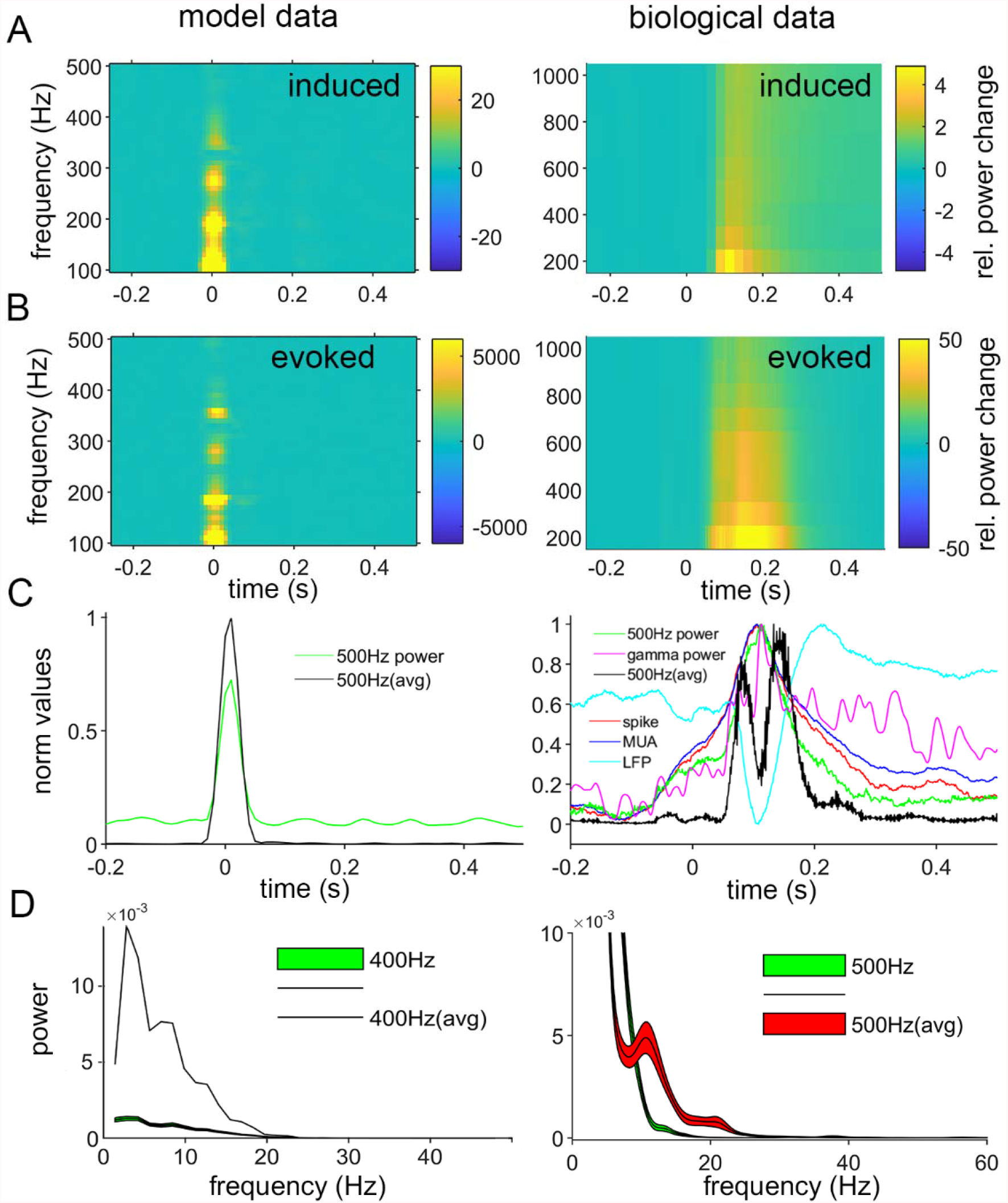
Comparison of model data from the area of stimulation to macaque V1 microelectrode recordings. Analysis of high frequency activity shown for data simulated for a small pick up area at the site of stimulation (top) compared to the microelectrode recording of V1 neurons responding to a visual stimulus (bottom). **(A)** Time frequency representation with respect to visual stimulus onset (time point 0). Induced (i.e. the mean over power values) broadband power increases are prevalent in the model data as well as the microelectrode recordings from macaque V1 (averaged over 20 sessions, 4863 trials in total). The relative power change refers to a baseline from -0.25 -0 s. **(B)** Time frequency representation of evoked (i.e. the frequency demodulation is applied after the time-domain average so only time-locked information is considered) broadband power increases. Otherwise same as in A. **(C)** The temporal evolution of the power (induced and evoked) in the 400 Hz (model) and 500 Hz band (biological data) (+/- 50 Hz, assessed in periods of 50 ms shifted in steps of 1 ms) is compared to spiking activity (summed over 50 ms, in steps of 1 ms), the MUA (absolute Hilbert transformed bandpass filtered 750 - 8000 Hz data), the LFP (lowpass filtered at 500 Hz) and the gamma power (FFT, 60 Hz). An exemplary session (178 trials) is plotted. **(D)** The evoked (red) and the induced (green) 400 Hz (model) and 500 Hz power change over time was frequency demodulated (FFT) to depict slow amplitude phase relationships. Only the evoked power shows a peak at 10 and 20 Hz. The colored area for the biological data depicts the SE over sessions.

**Figure 5:**
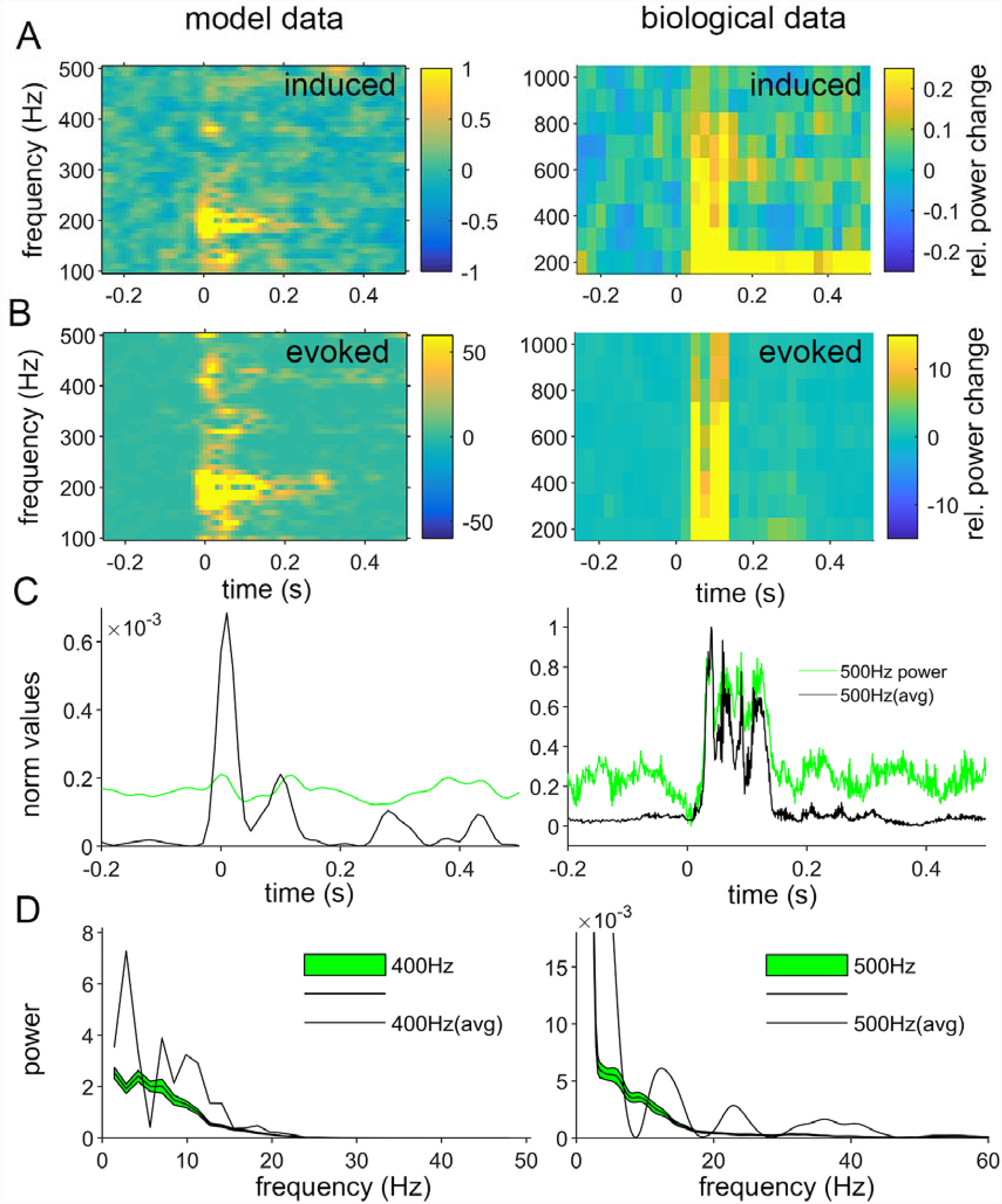
Comparison of simulation (data from an extended area) to macaque ECoG recordings. Analysis of high frequency activity shown for data simulated for a pick up area that includes the site of stimulation as well as neighboring sites (left) compared to the ECoG recording of V1 during visual stimulation (bottom). **(A)** Time frequency representation with respect to visual stimulus onset t (time point 0). Induced (mean over power values) broadband power increases are prevalent in the model data as well as ECoG recordings from macaque V1 (averaged over 73 trials). In the TFR, a single electrode above V1 is shown, (see the topographical representation and power distribution over the whole ECoG grid in **Fig S17C)**. The relative power change refers to a baseline from -0.25 -0 s. **(B)** Time frequency representation of evoked (i.e. the frequency demodulation is applied after the time-domain average so only time-locked information is considered) broadband power increases (see the topographical power distribution in **Fig S17D)**. Otherwise, same as in A. **(C)** The temporal evolution of the power in the 500 Hz (+/- 50 Hz, assessed in periods of 50 ms shifted in steps of 1 ms) band are compared between the evoked (green, 500Hz power) and induced signal. **(D)** The evoked (black) and the induced (green) 400 and 500 Hz power change was frequency demodulated (FFT) to depict slow amplitude phase relationships. Only the evoked power shows a slow modulation. No session-wise SE could be calculated.

## Discussion

We present a detailed open-source simulation for non-local information processing in a neocolumnar architecture and compare model output with *in vivo* neurophysiological data. Our work indicates that non-local information processing can be at the core of complex information coding. By this the same information is provided to all participating Mountcastle columns. With higher numbers of neurons involved, information integration by wave patterns emerges spontaneously as non-local information processing increases. Our modelling and experimental data on visual perception in an animal model support that high frequency neural activity encodes sensory information, which can be distributed via non-local, low frequency wave-like patterns across the cortex.

Previously applied large-scale models of the brain include the neocolumnar architecture ^2, 9^ and first efforts for multimodal neuroanatomic models ^27^. Furthermore, neuron simulations have also introduced new concepts such as aggregate-label learning ^28^. Our model on non-local information processing is generic and general, and just requiring a platform of microcircuits that are laterally interconnected. This can lead to shared information within cortical areas and inter-areal binding in a broad frequency range as information medium, and agrees well with observations from *in vivo* electrophysiological recordings (e.g. EEG or ECoG) including different pathologies. This non-local network architecture extends concepts of positive and negative feedback loops in cellular network architectures to a new emergent level. When cell networks are processing locally and modular, the non-local architecture allows for redundant copies of information and holistic distribution of information, so that each node in the network gets the same amount of information.

At the same time our model profits from and requires only a neocolumnar architecture, as present in the human brain ^2^. Information is encoded as a whole in time and space ^10^ thus forming interference pattern, that can emerge as clear waves. We focus here on the integrative properties of the model ^5^ at the basis of criticality ^15^ that indicate the maximization of information integration in frequency and phase. This processing platform serves as an intersection for continuous processing of world information in a positive and negative feedback loop ^29^.

Here, we demonstrate with our model that HF coding self-organizes at maximum frequency processing due to favored resonance bands of the model that is controlled by the energy coupling parameter. HF coding is only masked by the effect of undersampling (**fig. S16**). An additional emergent level is achieved by increasing the number of processing units, or neocolumns, that increase the stable resonances for frequencies (**fig. S11**). This adds growing phase information (**S19 and S20**), allowing unit by unit a more complex and stable representation of information. By potentiating frequency with phase, encoded complexity grows exponentially, and soon critical distributions arise that indicate systems that maximize information integration ^15^.

In the living system, were we have a sufficient amount of processing units (e.g. microcircuits, neocortical columns), high frequency activity of the brain is typically captured as multi-unit activity (MUA), a neural correlate of spiking activity ^30, 31^. HF activity shows an inter-areal phase coupling between task relevant areas in a visuo-motor task ^32^. Importantly, the here described evoked, time-locked HF changes that are observed during visual processing are distinct from MUA and LFP. They exist concurrently in and near the site of sensory processing. Only this evoked HF signal showed a slow phase (∼10 Hz) modulation. This novel observation suggests that the time-locked HF output is ordered by a slow phase pattern. This coupling between high and low frequencies might form a fundamental core of neural activity modulation and the coupling arises during sensory processing within a cortical area.

A prediction of our model applicable to the brain and its anatomy is that when a critical number of neurons in the brain is reached, a holographic medium might be able to integrate motor, proprioceptive and sensory input, to into a unified model of self and world representation. Mini-columns, conceptually part of cortical columns ^33^ contain about 80-100 neurons ^9^ and about 50 - 100 minicolumns are organized in a cortical column ^33^. In our simulation, about 14,400 neocortical columns with about 10^4^ neurons per column ^2^ allow sufficient resolution to store accurate wave patterns. This gives a very rough estimate regarding the theoretical lower limit required for emergence of such non-local patterns in brain areas, like the visual cortex. Our model predicts that only a sufficiently high number of neurons organized in a non-local architecture allows to maximize information integration. This might be a prerequisite for integrating sufficient information to ultimately reach consciousness.

In summary, simulations and collected observational data all support our central hypothesis of non-local, wave-like processing of information in the cortex as a root-phenomenon for higher brain functions. Here we transfer non-local information processing requiring just a columnar architecture. Like the higher primate cortex, the neopallium of birds has been proven to be suited for processing of perceptual and cognitive abilities and recently, it was found to have a specific columnar architecture ^34, 35^ which, according to our computer model, should be similarly well adapted to non-local information processing. Such convergent evolution in different organism groups (mammals, birds, and maybe others) is a striking argument that the properties of a columnar architecture are important for higher brain function.

## Acknowledgements

We thank Pascal Fries for allowing to reanalyze already published macaque ECoG data and Thilo Womelsdorf for sharing unpublished macaque microelectrode data with the authors.

## Author contributions

JB set-up, tested, analyzed and finalized the brain simulation including data comparisons to experiments and was supervised by TD. BH enabled the use of the animal data and analyzed all animal recordings as well as model data used for comparison. JP set-up and did the protein interaction circuit simulations within a neuron. RB generated and provided complementary neuronal oscillation data and videos. BH, CAB, HE, SMW, RB and TD provided neurobiological expertise. JG and JvK supervised by SK did large-scale grid computing simulations on the brain simulation code and parallelized it with input from JB. SMW provided MEA data and did the connected experiments on hippocampal brain slices. RB and TD led and guided the study. JB, RB, TD drafted the original manuscript. All authors (JB, BH, JP, CAB, HE, JG, JvK, SMW, SK, RB, TD) edited the manuscript, gave comments and agreed to its final version.

## Data and materials availability statement

All data for this study are contained in the manuscript, its figures and the supplements. This includes links to download the complete used program code and a tutorial for its use. Also the code used to process the animal data and to obtain the shown data figures is made fully available. We allow for data redistribution for the purpose of replication.

## Supplementary materials

Supplementary document containing extended methods, extended results, and extended discussion; table S1, S2, supplementary figures S1-S20. Independent files are table S1 (excel file Trk receptor protein-protein interaction network) and fig S3 (high resolution figure on Trk interaction network and Jimena analysis). Video material: video S1-S13.

