## Supplemental Material for "Modelling non-local neural information processing in the brain"

**Content**

### Extended figure legends (main text - Figures 1 – 5)

Extended legend - Figure 1: A non-local information processing model is embedded in a neocolumnar architecture. **(A)** In order to simulate non-local information processing, three processing steps, a spatial derivation and two temporal integrations, built on each other to allow simulation and interconnections of microcircuits and their parallelizability. (see **eq. 1**): First, the derivation of the neighboring energy (*E_N(t-1);* *activation1_n_(t)*) and the own energy state (*E(t-1); activation1(t)*) leaded to the *Delta’(t)* value (*slope_vector(t)*) of that specific node. Second, this *Delta’(t)* value was integrated with the *Delta(t-1)* function (*slope_old(t-1)*). Third, the updated *Delta(t)* (*slope_old(t)*) was added to the energy state function of the focused node (*E(t-1); activation1(t)*) leading to the current final energy state (*E(t); activation0(t)*). Both the new delta (*Delta(t)*; *slope_old(t)*) and the new energy state (*E(t); activation0(t)*) were then forwarded to the next time step, while the current energy state (*E(t-1);activation1(t)*) was forwarded to all neighboring nodes. The variables *a* (*ratio_neighbor_activation*), *b* (*ratio_inhibition_activation1*), *c* (*neighbor_integration; NI_slopev*), *d, e, f* (*slopeo_damping*) and g (*damping*) were coupling factors. These modulated the energy transfer between the neuronal nodes. Their default value is 1. The variables can be replaced by a function, e.g. a damping function. **(B)** A computer cluster simulation of non-local processing. Shown is a perception task model with 5,000 ticks with a resolution of 1 tick per millisecond (i.e., 5 seconds of real-time) and its dependence on the number of simulated nodes in the cluster. The perception task was done more efficiently with more nodes but there were no emergent new properties.

Parameters: Average run-time of 5 repetitions are shown as lines, the minimum and the maximum run-time as semi-opaque areas of the same color. We were able to simulate up to 50,000 nodes in real-time (i.e., the simulation runs faster than the modelled process itself) using a standard 2-core laptop. However, it was possible to simulate even more nodes (100,000 and higher) by running the simulation on server CPUs. We furthermore observed that the servers show less random influences compared to the laptop workstation. This was expected, as we used bare-metal servers with no other competing processes running at the time of the simulation. Additionally, we observed that both the laptop and the smaller Skylake server were faster for smaller simulations but were strongly influenced by the number of nodes to simulate. The reason for this is the smaller L2-cache. In fact, we observed a strong impact on the performance of the Skylake server at around 500,000 nodes. This was exactly when the node grid got too big for the L2-cache, which had a serious impact on performance. **(C)** A neuronal circuit for non-local information processing. The holographic brain circuit allows processing calculations, high discrimination, and signal resolution (compare **eq. 1** and **(A)**). The innercolumnar network was represented by neuronal microcircuits and its connections to the neighboring neurons (*activation1 N1-4*). An inhibitory neuron (*activation0/1* in red) stores the previous energy state and subtracts it from the neighboring excitatory neuron energies (*activation1 N1-4*). This current energy derivative (*slope_vector*) is updated on the excitatory neurons *slope_old* that integrates its new energy level on an excitatory (*activation0/1* in blue; external transmission) and an inhibitory neuron (*activation0/1* in red; internal feedback). The energy transmission can be modulated by ion channels, such as GABA receptors, that dampen the cell body energy levels (*damping* and *slopeo_damping*) or dampen near synaptic energy transmission (*NI_slopev*). A complex interference pattern was formed when processing stimuli on a platform of interconnected microcircuits. Input signals are interfered and integrated existing information. As a consequence, the entered signals were distributed over the whole model, which made it available at any column after a distinct time period. Processing of information was dependent on time and space. Thus, the information could be coded in frequency and phase. With a Fourier transform the signals were analyzed. Here, complex input to the model was defined by more than 50 stimuli per second with a duration of 50-100 ms and an information bandwidth of 200-500 Hz superimposed on reference signal of 1-20 Hz. Small electrodes could record bursts of fast ripple-like signals. If measuring the same spot at the same time, signals from big electrodes appear similar to the topology of event related potentials. The model could represent a platform for interfering afferent stimuli (e.g. sensory input). Afferent stimuli, like motor actions can be deduced from the subsequent interference pattern at every location of the model. Thus, the model could represent a continuous information feedback model.

Extended legend - Figure 2: Analysis of the neocolumnar non-local information processing simulation. Distinct characteristics of the simulations are rooted by the energy transmission of the lateral interconnections. Fundamental energy features are also found in electrophysiological recordings of the brain and can be utilized to discriminate brain states. **(A)** The modelled waking state is indicated by low in amplitude, fast oscillations. A decrease of the energy coupling parameter *NI_slopev* from 2.6655 to 0.1 induced the slow wave state (SWS), which was characterized by slow oscillations that were large in amplitude. Further, a slight decrease of *damping* (1.0E-4🡪1.0E-5), *slopeo_damping* (1.0E-2🡪1.0E-3) and the reference signal frequency (0-23🡪0-13 Hz) facilitated the smooth low frequency oscillations in SWS state. Further simulations were compared to the waking state parameter space. The size distribution of positive peaks reproduced MEA peak distribution in **(B)** waking and SWS state and **(C)** anesthesia state (*damping* and *slopeo_damping* were 1.0E-3 and 1.0E-1, whereas *NI_slopev* stayed at 2.6655). Waking and SWS state signals were lognormal distributed, whereas anesthesia state signals were power law distributed. (**D**) Similar lognormal distributions were generated by electrode recordings of hippocampal brain slices of a 33 day old wt mouse. Shown are the baseline activity for 15 min and periods after stimulation with CCH (carbachol) at 0-15, 15-30, 30-45 and 45-60 min (detailed analysis in **fig. S6**). **(E)** Systematic analysis of harmonics generated by the simulation. The input was a continuous periodic peak signal of 7 Hz.

###### Figure2 – model parameters

**Fig. 2A:** EEG signal: 21 sine-like stimuli of 400-500 Hz with 800mV amplitude and a random onset of bursts (average 250 ms) of 50 ms length were applied. 20% of stimuli were superimposed on a reference signal 0-30 Hz with 2400 mV strength. The input of slow wave state and waking state were alike despite an increase of *NI_slopev* from 0.1 to 2.6655 accompanied by a slight increase of *slopeo_damping* (1.0E-5🡪1.0E-4) and *damping* (1.0E-3🡪1.0E-2) decreases the amplitudes of waves and facilitates gamma oscillations. Raising the carrier frequency from 0-13 Hz to 0-23 Hz favored the usage of the resonance frequency (compare baseline in **Fig. 3C** and **S12**). In the slow wave state (SWS), low carrier frequencies enhanced the effect of amplitude increase. The decrease of *NI_slopev* functioned like a low pass filter. The EEG-electrode radius was 10 mm. The *marginextradamping* was 2.

**Fig. 2B, C:** The positive peak distribution of the simulated MEA signals for SWS and waking state are indicated in **Fig. 2B**. The according parameters are described in **Fig. 2A**. Turning from waking state to anesthesia urged an increase of *slopeo_damping* (1.0E-2 🡪 1.0E-1) and *damping* (1.0E-4 🡪 1.0E-3). The *NI_slopev* (2.6655) in anesthesia was the same as in waking state.

**Fig. 2E**: The generation of harmonics used 7 Hz continuous peak input, a *NI_slopev* of 2.6655, a *slopeo_damping* of 0.01, a *damping* of 0.0001, a *marginextradamping* of 2, an *inputstrength* of 800 mV and a simulation length of 3 s. The measurements were done on LFP recordings, which were simply generated by an electrode size (radius = 1 mm).

Extended legend - Figure 3: Simulation and modulation of neocolumnar non-local processing of stimuli. (**A**) Efficient resolution of 28 different stimuli as complex model input. The input was coded in prime numbers. The corresponding frequencies were 5 - 250 Hz. 28 out of 29 input signals and 45 harmonics could be decoded. For comparison, the undetected frequency was in the LF space (<7 Hz). The additional decoding of harmonics implied that the model can easily encode more than 50 stimuli. **(B)** The favored frequency bands of the model are analyzed when simulated spontaneous activity only is applied (0.01% and input strength of 100 mV). While waking showed a favored bandwidth of >300 Hz in the MEA signal, SWS showed a favored bandwidth of <100 Hz. The decrease of *NI_slopev* from >2.6 to 0.1 indicated a low pass filter effect. Anesthesia decreased the energy in all frequency bands. **(C)** Self-organizing baseline activity of the simulation mimicked observations. The baseline organized after applying spontaneous activity only (100 mV and 0.01% activated columns). **(D)** Coherence declines with frequency and space in the model. MEA coherence was measured in presence of spontaneous activity (0.01% activated cells firing with 100 mV). The simulation length was 10 seconds. The distance of the MEA electrodes was 0.5 (MEA1-2), 2 (MEA1-2), 4 (MEA1-3), 8 (MEA1-4), 16 (MEA1-5), 32 mm (MEA6). (**E)** The Lempel Ziv complexity of the EEG recordings is giving hints about the complexity of information processing at different modeled brains states. LZC of brain recordings varied between ordered sine waves (LZC=0.2) and gaussian noise (LZC=1.3). With LZC brain states, such as waking, rapid-eye-movement sleep (REM), slow wave sleep (SWS), AD, schizophrenia, Ketamin and Propofol anesthesia could be differentiated. With LZC the modeled states waking (**Fig. 3A**), SWS (**Fig. 3A**), REM sleep (change waking *NI_slopev* 2.6655🡪1), AD with 1 % and 10 % of lesions (**Fig. 4F**), schizophrenia (**Fig. 3G+H**), Ketamin-induced anesthesia (**Fig. 3C**) and Propofol-induced anesthesia (Ketamin anesthesia with a shift of *NI_slopev* 2.6655->0.1) were anaylzed. **(F)** Influence of lesions occurring during Alzheimer’s disease on information processing. Axis and parameters were similar to **fig. S10**, except the model size was fixed (120 x 120 columns) and the x-axis represents the ratio of lesions. The left y-axis shows the number of harmonics and artefacts. The right y-axis shows the number of matches. The matches are the frequencies of the output of the simulation that matched to the input frequencies. The harmonics could be deduced from input frequencies. The artefacts had no clear relation to the input frequencies but were clear frequencies in frequency space. **(G)** Low correlation such as in schizophrenia decreases information content and input separation. Axis and parameters are as in **(F)**. The percentage of connections between columns showing uncorrelated energy transfer is indicated by the x-axis. **(H)** **Gamma-band decline in schizophrenia.** A self-organized gamma-band superimposed on a reference wave followed a complex stimulus as shown for healthy (left) subjects and schizophrenic subjects (middle). The delta of the signals of the healthy model (left) and the signal of the schizophrenia model (middle) in frequency space is shown in the (right) panel. The model parameters (left) were the same as in **Fig. 2A** (simulated waking state). However, the reference signal frequency was fixed to 10 Hz and the superimposed HF stimulus was fixed to 430 Hz. The burst length was 50 ms and burst onset was fixed to 2 s after simulation start. The color bar indicates the activation in µV.

###### Figure 3 – model parameters

**Fig. 3A**: The parameters are follows: 29 continuous peak prime inputs of 5-239 Hz, *NI_slopev* = 2.6655, *slope_damping* = 0.01, *damping* = 0.0001, *marginextradamping* = 2, signal length = 3000 ms. The signals were recorded by simulated MEA signals.

**Fig. 3B**: The parameter for waking, SWS were as in Fig. 2A and for anesthesia as in Fig. 2C. The windows size for the running average was 120 *ms* and the signal length was 9 *s*. The simulated MEA signal was analyzed.

**Fig. 3C**: Baseline is generated by applying a virtual spontaneous activity of 100 mV and 0.01 % of randomly activated columns each time step. Further parameters were as follows: *NI_slopev* = 2.6655, *slopeo_damping* = 1.0E-2. *damping* = 1.0E-4, *marginextradamping* = 2.0 and signal length = 10000 ms. Here, the simulated EEG signals were analyzed (diameter = 2 cm).

**Fig. 3D**: The coherence was measured by a simulated MEA. The input for the simulation wa spontaneous activity (0.01 % activated cells firing with 100 *mV*). The signal length was 10 s. The *NI_slopev* was 2.6655, *slopeo_damping* was 1.0E-2, the *damping* was 0.0001 and *marginextradamping* was 2.0. The distance of the MEA electrodes was 0.5 (MEA1-2), 2 (MEA1-2), 4 (MEA1-3), 8 (MEA1-4), 16 (MEA1-5), 32 mm (MEA6).

**Fig. 3E**: The Lempel Ziv complexity was estimated for the simulated EEG signals (radius = 2 cm). The parameter for the simulated waking state and SWS state were the same as in Fig. 2A. The REM state has the same parameter values as the waking state but the *NI_slopev* is decreased from 2.6655 to 1. Two disease states, AD (parameter explained in Fig. 3F: 1 % and 10 % lesions) and schizophrenia (parameter explained in Fig. 3G: 1.0 % uncorrelated), as well as two modeled states of anesthesia (Ketamin and Propofol) were simulated. The modeled Ketamin-induced anesthesia has the same parameter values as waking but the *slopeo_old* was set to 1.0E-1 and the *damping* was set to 1.0E-3. The modeled Propofol anesthesia has the same parameter values as the Ketamin-induced anesthesia but the *NI_slopev* was additionally decreased to 1.0E-1. The simulated EEG signal was analyzed for 8 s. The signal was transferred into a binary code by using the median of the signal period. The Lempel Ziv algorithm of Matlab R2020a was used for calculating the complexity value.

**Fig. 3F,G**: The simulation of AD and schizophrenia are based on the same parameter values as the simulated waking state shown in Fig. 2A. In difference to a simulated healthy model (here waking) the AD model introduces lesion (the x-axis showed percentage of lesions). In schizophrenia the x-axis showed the percentage of connection with randomly declined energy transfer (percentage uncorrelated). The simulation was equally resolving 12 continuous peak stimuli (29, 31 37, 41, 43, 47, 53, 59, 61, 67, 71, 73 Hz) in the healthy state (waking). The matched frequencies, harmonics, and artificial frequencies were investigated. The left y-axis corresponds to the number of detected artificial signals and harmonics. The right y-axis represents the number of input frequencies. Further parameters: *NI_slopev* = 2.6655, *slopeo_damping* = 0.01. *damping* = 0.0001, *marginextradamping* = 2.0 and signal length = 3 s. The simulated MEA signals were analyzed.

**Fig. 3G**: For the gamma-band decline analysis in simulated schizophrenia, the input was of sinus wave type. Information was transmitted in a burst of 50 ms. The information was superimposed on a reference signal of 10 Hz and the information was coded in HF space (>400 Hz). The energy coupling of the columns was randomly decreased to values between 1 and 0.8 in 99 % of the columns. The regarding model parameter were *uncorrelated4_slopeo*, *uncorrelated* and *uncorrelated_input*. The parameter *uncorrelated3_slopev* enabled to randomly decreased the *NI_slopev* to values between 1 and 0.8 in 12.5 % of the connections between the columns. Further parameters: *NI_slopev* = 2.6655, *slopeo_damping* = 0.01, *damping* = 0.0001, *marginextradamping* = 2.0 and signal length = 10,000 ms. The simulated EEG signals were analyzed using a radius of 1 cm.

Extended legend - Figure 4**: Comparison of model data from the area of stimulation to macaque V1 microelectrode recordings.** Analysis of high frequency activity shown for data simulated for a small pick up area at the site of stimulation compared to the microelectrode recording of V1 neurons responding to a visual stimulus (bottom). **(A)** Time frequency representation with respect to visual stimulus onset (time point 0). Induced (i.e. the mean over power values) broadband power increases are prevalent in the model data as well as the microelectrode recordings from macaque V1 (averaged over 20 sessions, 4863 trials in total). The relative power change refers to a baseline from -0.25 -0 s. **(B)** Time frequency representation of evoked (i.e. the frequency demodulation is applied after the time-domain average so only time-locked information is considered) broadband power increases. Otherwise, same as in A. **(C)** The temporal evolution of the power (induced and evoked) in the 400 Hz (model) and 500 Hz band (biological data) (+/- 50 Hz, assessed in periods of 50 ms shifted in steps of 1 ms) is compared to spiking activity (summed over 50 ms, in steps of 1 ms), the MUA (absolute Hilbert transformed bandpass filtered 750 - 8000 Hz data), the LFP (lowpass filtered at 500 Hz) and the gamma power (FFT, 60 Hz). An exemplary session (178 trials) is plotted. **(D)** The evoked (red) and the induced (green) 400 Hz (model) and 500 Hz power change over time was frequency demodulated (FFT) to depict slow amplitude phase relationships. Only the evoked power shows a peak at 10 and 20 Hz. The colored area for the biological data depicts the SE over sessions.

Extended legend - Figure 5**: Comparison of model data from an extended area to macaque ECoG recordings.** Analysis of high frequency activity shown for data simulated for a pick-up area that includes the site of stimulation as well as neighboring sites (top) compared to the ECoG recording of V1 during visual stimulation (bottom). **(A)** Time frequency representation with respect to visual stimulus onset t (time point 0). Induced (mean over power values) broadband power increases are prevalent in the model data as well as ECoG recordings from macaque V1 (averaged over 73 trials). In the TFR, a single electrode above V1 is shown, (see the topographical representation and power distribution over the whole ECoG grid in **Fig S17C)**. The relative power change refers to a baseline from -0.25 -0 s. **(B)** Time frequency representation of evoked (i.e. the frequency demodulation is applied after the time-domain average so only time-locked information is considered) broadband power increases (see the topographical power distribution in **Fig S17D).** Otherwise, same as in A. **(C)** The temporal evolution of the power in the 500 Hz (+/- 50 Hz, assessed in periods of 50 ms shifted in steps of 1 ms) band are compared between the evoked (green, 500Hz power) and induced signal. **(D)** The evoked (black) and the induced (green) 400 and 500 Hz power change was frequency demodulated (FFT) to depict slow amplitude phase relationships. Only the evoked power shows a slow modulation. No session-wise SE could be calculated.

### Supplementary Figures

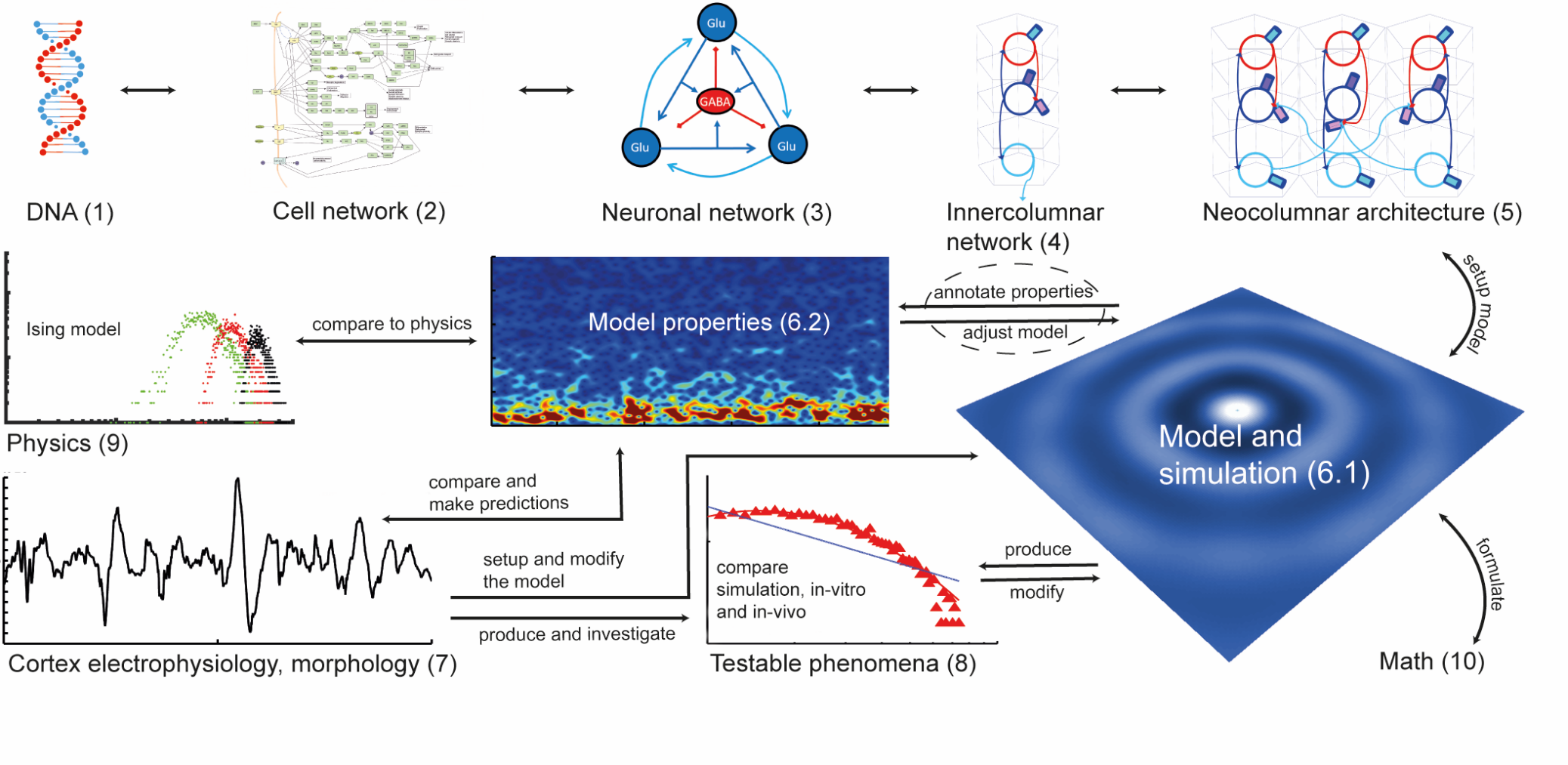

Fig. S1: Workflow for investigating non-local information processing in cortical structures.

This study models non-local processing in a neocolumnar, cortex-like architecture. This figure describes basic considerations of how cortical information is generated on different levels. The architecture of the cortex is stored on the DNA (1) which gives rise to cellular networks and thus build up internal and external cellular communication pathways (2). Neurons use cells as intersection in a neuronal network and (3) are organized in the cortex in columnar structures, functioning as a complex processing unit (4). These neocortical columns are again interconnected building up neocortical architecture (5). We believe that this architecture forms a platform for non-local information processing. Simplifying the rules of communication of the columns enables a transfer to an *in silico* model and a subsequent simulation of this architecture (6.1). The processing steps of the simulation are closely adopted to biological parameter, as well as the input signals resemble biological signals. Thereby model properties are categorized (6.2) and can be compared with biological data, for instance with electrophysiological data acquired from cortical structures (e.g. the visual cortex, see Fig. 4, 5) (7). Phenomena that are investigated, analyzed and described by *in vivo* and *in vitro* studies are reproduced by the model (8). The comparison of *in silico* and *in vitro*, or *in vivo* data can be used for adjusting the model parameters and dynamics. Using this feedback loop the simulation can become more and more biological. A similar feedback approach compares the simulation with physical models (9), as well as mathematical formulas and phenomena (10).

In summary, we have set up a neocortical model that implements principles of mathematics, physics, and biology. The model combines robust information processing properties with usability to achieve annotation, description, comparison, and simulation of physiological phenomena of higher brain functions.

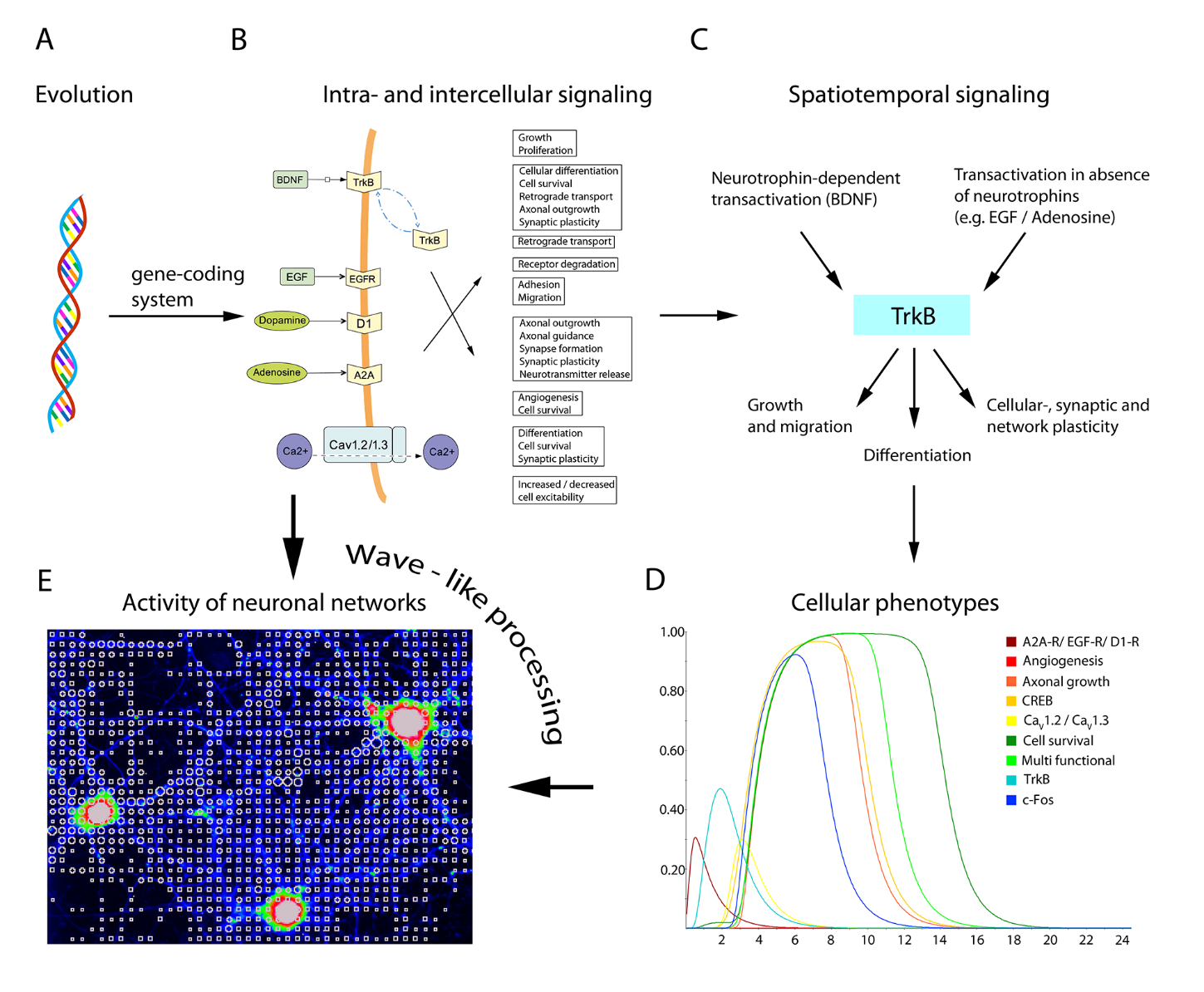

Figure S2: Information is processed on ever higher levels and networks. Evolution selects information for better and better adaptation and survival. Information is stored internally first in DNA, but then this is combined with ever higher levels of nodes (proteins, neurons …) and super-nodes (protein networks, neocolumns…). This process chain is sufficient to lead to rather complex emergent behavior. Different emergent levels of signal processing are exemplarily illustrated to show how wave-like processing is based on simple rules, but contributes to emergence. **(A)** Level 1 – Evolution: Genetic information codes the regulation program of cell type-specific signaling networks. **(B)** Level 2 – Signaling (according to a representative signaling cascade): Signaling caused by the brain-derived neurotrophic factor (BDNF) stimulates cellular and synaptic plasticity in cortical regions of the brain, e.g. by activating the cell surface TrkB and intracellular TrkB (two blue arrows, top) embedded in a complex intra- and intercellular signaling system. **(C)** Level 3 – spatiotemporal signaling (according to a representative signaling aspect): Spatiotemporal activation of diverse cascades is caused by neurotrophin-dependent effects through BDNF/TrkB or by transactivation of TrkB in the absence of neurotrophins for migration, differentiation, or survival of neural cells in development, differentiation, and plasticity in the adult brain. **(D,** right**)** Level 4 - Signaling interactions between cells: neurotrophin-independent activation of TrkB by either adenosine receptor A2A-R, EGF-receptor (EGF-R), or dopamine receptor D1 (D1-R) activates specific key nodes (e.g. the transcription factor CREB) for different cellular functions, such as information processing by regulated cell surface abundance of TrkB **(E,** left**)**. Level 5 – emergence level: Information, as those described on level 1 to 4 is a fundament for physiological signal integration, as for instance seen in synchronized neuronal activity with wave-like properties, e.g. by chemical activation of hippocampal neurons in culture (**suppl. video S1** and **S2**).

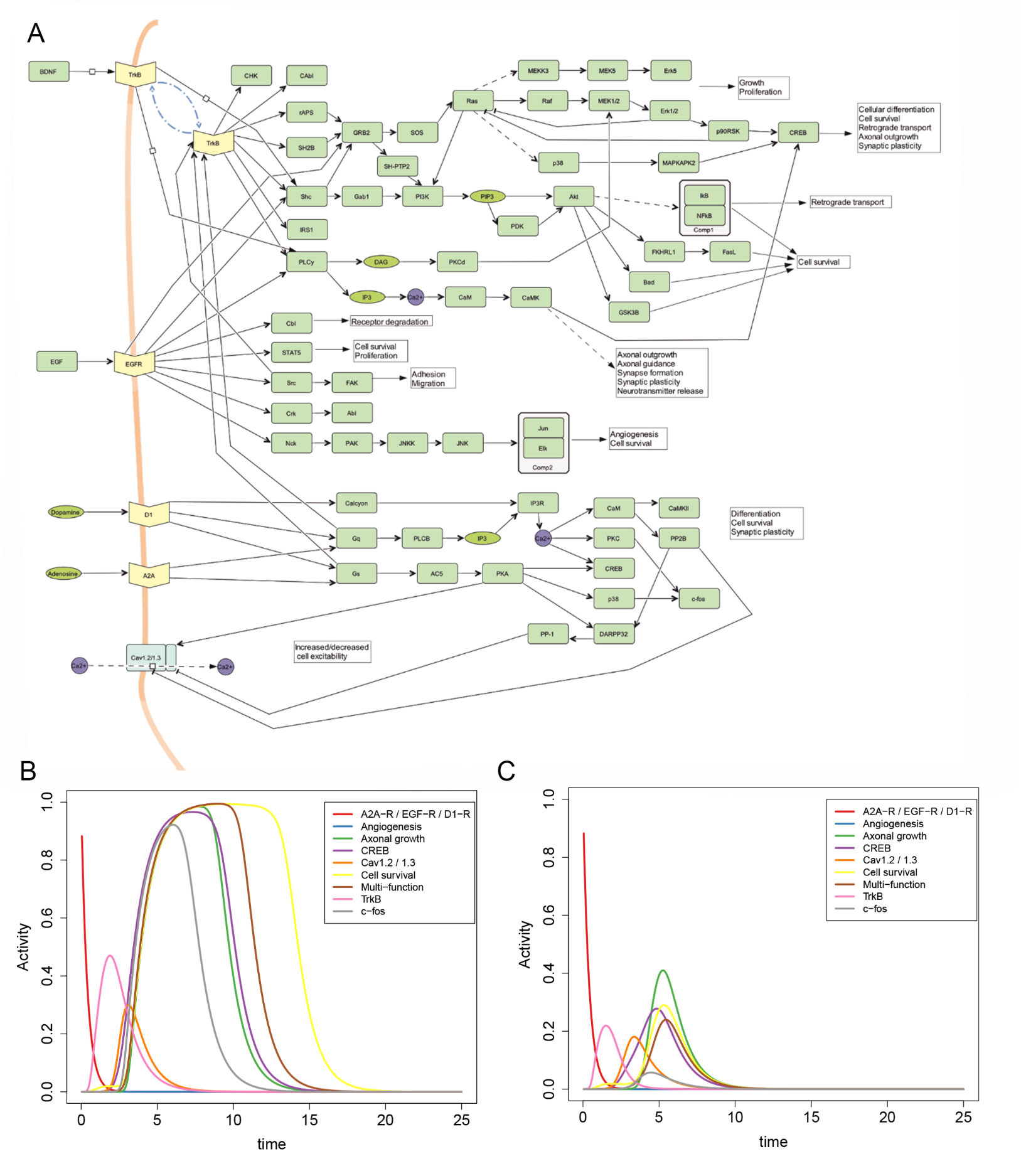

Fig. S3: Cellular model of the BDNF / TrkB signaling network (electronic figure version)**: (A)** This model (protein network, level 2) is presented in **Fig. S2B and C** and is given here in detail. The proteins for this are encoded on the DNA level (basic level 1). It is based on **Table S1** (separate Table in excel format; all nodes, interactions, references; hand curated) and was assembled starting from the KEGG pathway “ko04722 Neurotrophin signaling pathway” ^1, 2^, the resulting network is shown in the separate file GraphS1.graphml in graphml format. This pathway presents the direct activation mechanisms of the neurotrophins NGF, BDNF, NT3 and NT4 and the subsequent activation of the intracellular signaling cascades including the pathways represented by MAPK, PI-3 kinase and PLC. The activation of these cascades results in functions like axonal growth, cell migration, cell survival, cellular differentiation and plasticity. The pathway was later modified to allow for different neurotrophin-independent TrkB transactivation routes. A first transactivation route starts with charged ions like zinc or calcium ^3^ passing through channels like VDCC or NMDAR. Another starting factor of TrkB transactivation is dopamine ^4^, which activates G-protein-coupled receptors and subsequently transactivates TrkB receptors. A third TrkB transactivation route is mediated by adenosine and the A2A adenosine receptor, another G-protein-coupled receptor ^5^. Finally, the growth factor EGF is also present as a TrkB transactivation factor ^6^. **(B) and (C): Cellular phenotypes including differentiation arise from modulating protein-protein interactions between cell compartments.** An extended version of **fig. S2D** is shown in **B** and **C.** The simulation illustrates how BDNF signaling versus neurotrophin-independent TrkB transactivation can result in different cellular responses and thereby affect multi-functional signaling cascades, but also functions such as axonal growth or cell survival. Important key factors such as the activity-dependent transcription factors CREB or c-FOS are pointed out. The simulation shows different results depending on whether neurotrophin-independent TrkB transactivation routes induced by the adenosine receptor A2A (A2A-R), EGF-receptor, or dopamine receptor D1-R are ON **(B)** or OFF **(C)**. Note that neurotrophin-independent TrkB activation can occur at intracellular sites and defined cellular functions no longer emerge when the decay factor of TrkB activation is changed. The neuronal protein-protein network simulation was run using the topology of the BDNF signaling network shown in **(A)** and its dynamic behavior is in line with experimental data. For instance, A2A-R-dependent activation of TrkB can contribute to neuronal survival, at least in motoneurons and blocking this transactivation route can reduce the A2A-R mediated effects ^5^.

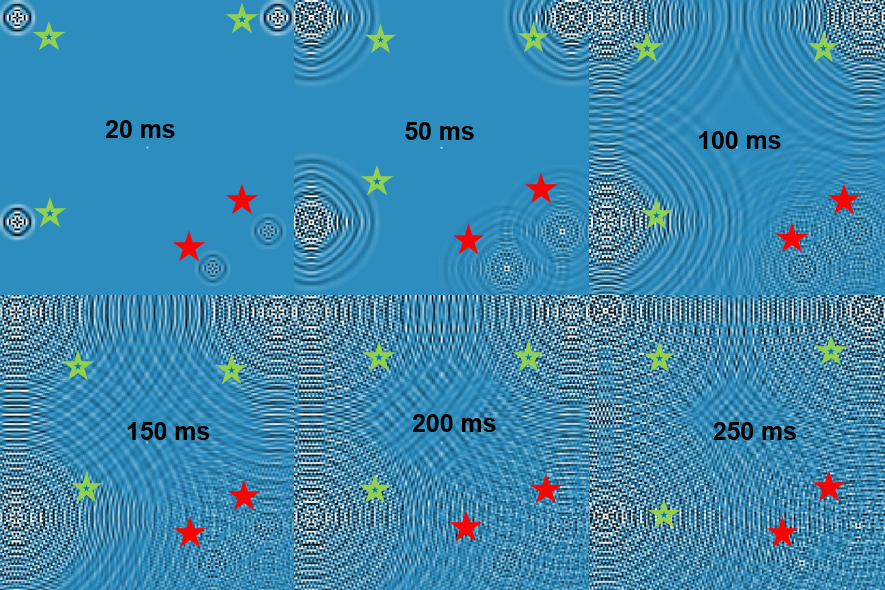

Figure S4: Waves arise at stimuli location, bump into each other after time and generate a complex interference pattern: Six screenshots show the development of three internal stimuli (green stars) and two external stimuli (red stars) in the non-local simulation at different time points (20, 50, 100, 150, 200 and 250 ms). It is shown that stimuli are transformed into a wave-like signal and that the interference pattern gets more complex with time.

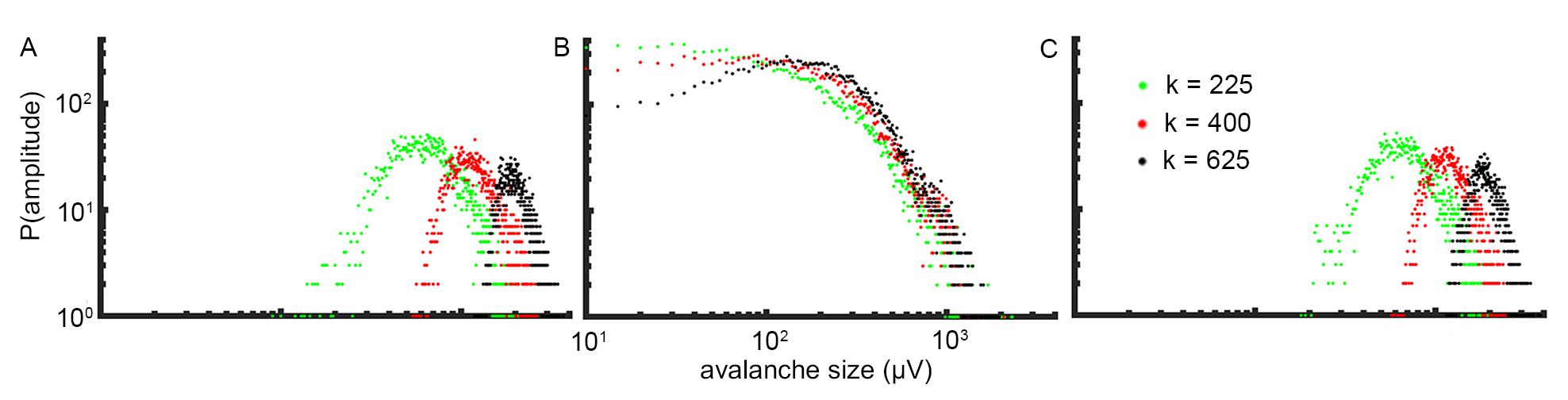

Figure S5: critical state of high information present in the non-local network simulation. The analysis of Ising-like dynamics in the simulation indicate that the critical state of high information integration can be stabilized and maintained. The parameter tuning indicates that the critical state in the non-local information processing simulation is very robust over a large parameter shift. For testing the relation of criticality to order and chaos in this model the *NI_slopev* was lowered to 5.0E-1. The variables *slopeo_damping* and *damping* were set to 5.0E-2 and 0 respectively. No margin was applied. The EEG radius is set to 2 mm. 0.02 % of the processing units were activated randomly each ms. In this variable space the model is still showing critical distribution **(B)**. The shift of values is necessary, as the values used for waking (compare to parameter space for maximum processing in waking state in **Fig 2B**: *NI_slopev* is 2.6655, *damping* and *slopeo_damping* is 1.0E-3 and 1.0E-1) are to robust for a shift to order and chaos. A further decrease of the *NI_slopev* to 5.0E-2 resulted in ordered processing. **(A)**. An increase *slopeo_damping* to 0.5 resulted in chaotic processing **(C)**. The distributions are generated for different model size with 225, 400 and 625 processing units. The model size is represented by the variable *<k>* which is used analog to the average network connectivity value in Fraiman et al. ^7^. An increased number of processing units connected within a network shifted the distribution to higher energies, which is valid for order, criticality, and chaos.

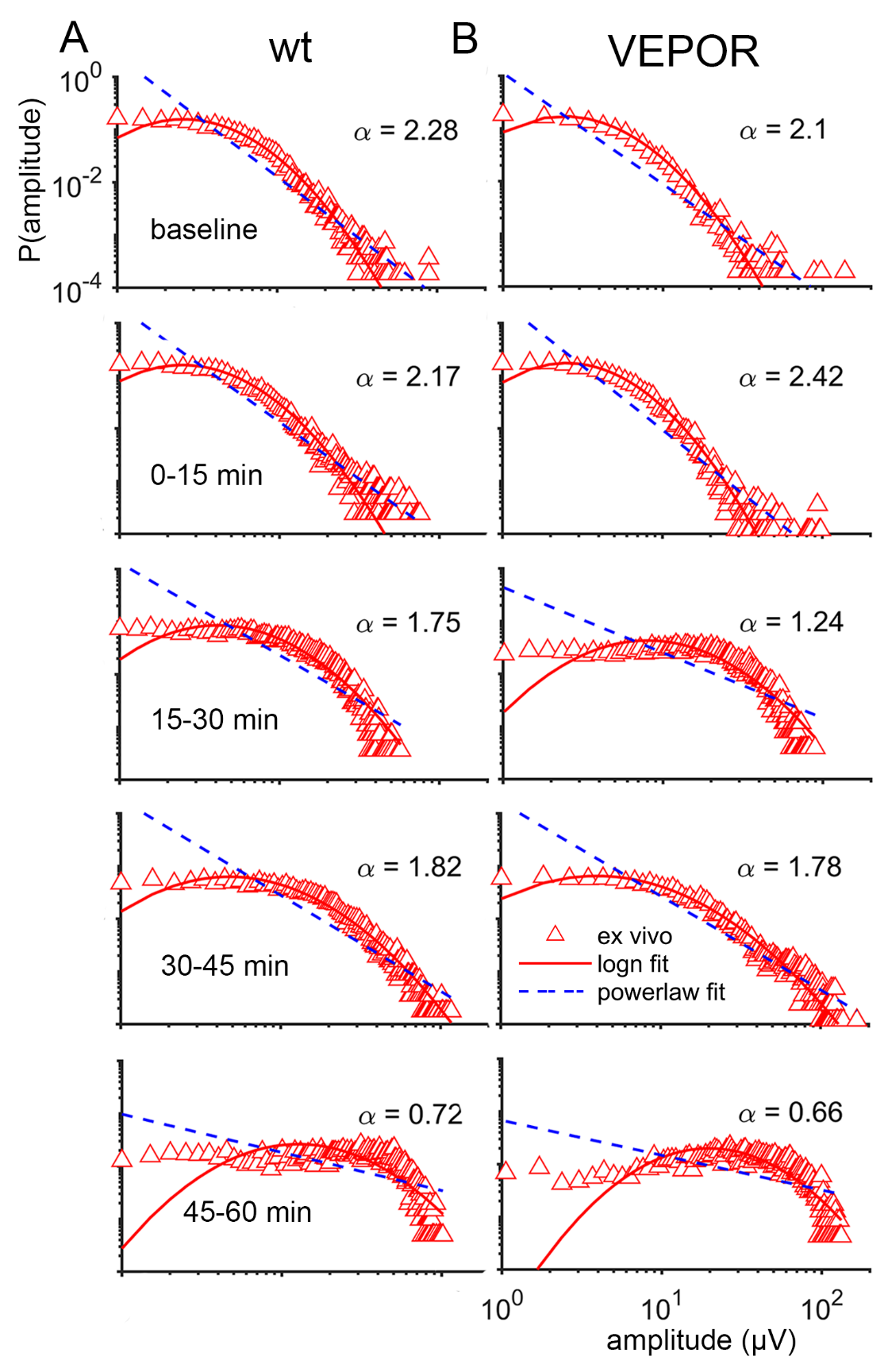

Figure S6: The analysis of the peak distribution of electrophysiological recordings of hippocampal brain slices indicates lognormal distributions. (**A**) We show here electrode recordings of hippocampal brains slice of a 33 day old wildtype mouse. Shown are the baseline activity for 15 min and periods after stimulation of acetylcholine receptors with the agonist carbachol (CCH) for 0-15, 15-30, 30-45, and 45-60 min. CCH seems to trigger shifts in the peak distribution, which might indicate altered information processing, or synchronized oscillations. (**B**) The electrode recordings of hippocampal brains slice of a 27 day old mouse with a constitutively active form of the erythropoietin receptor in GABAergic neurons (VEPOR+/+) ^8^. As in (**A**) the baseline activity for 15 min and periods after stimulation with CCH for 0-15, 15-30, 30-45 and 45-60 min are demonstrated. Comparing (**A**) and (**B**) shows a qualitatively similar shift in the signal peak distribution induced by CCH in wildtype as well as the genetically modified VEPOR mice.

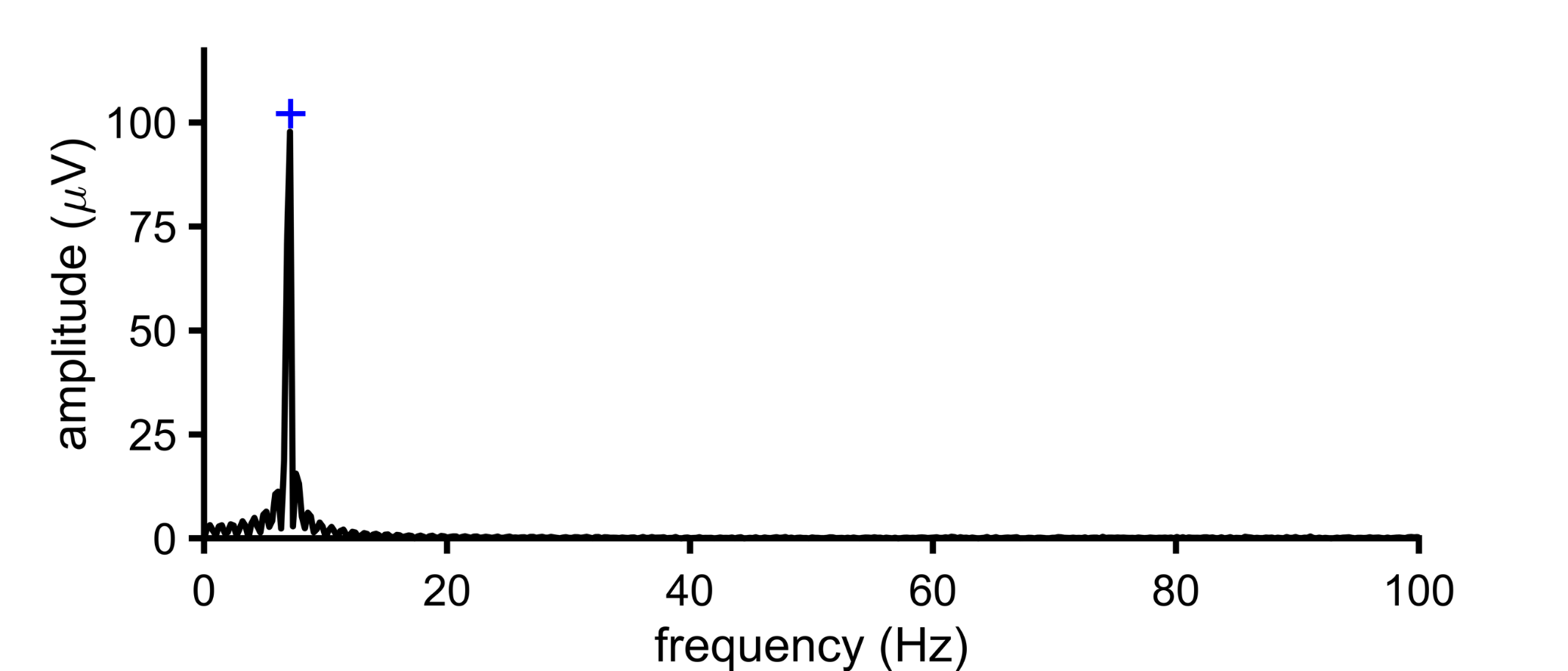

Figure S7: Sinus rhythm extinguishes harmonics. Shown is the analysis of harmonic decline in presence of sine input generated by the simulation. For demonstrating harmonic decline, a 7 Hz continuous sine input was applied. Other parameters: *NI_slopev* of 2.6655, *slopeo_damping* of 0.01, *damping* of 0.0001, *marginextradamping* of 2, *inputstrength* of 800 mV and a simulation length of 3 s. The simulated local field potential (LFP) signal was defined as the summed activity of simulated columns within a radius of 1 mm.

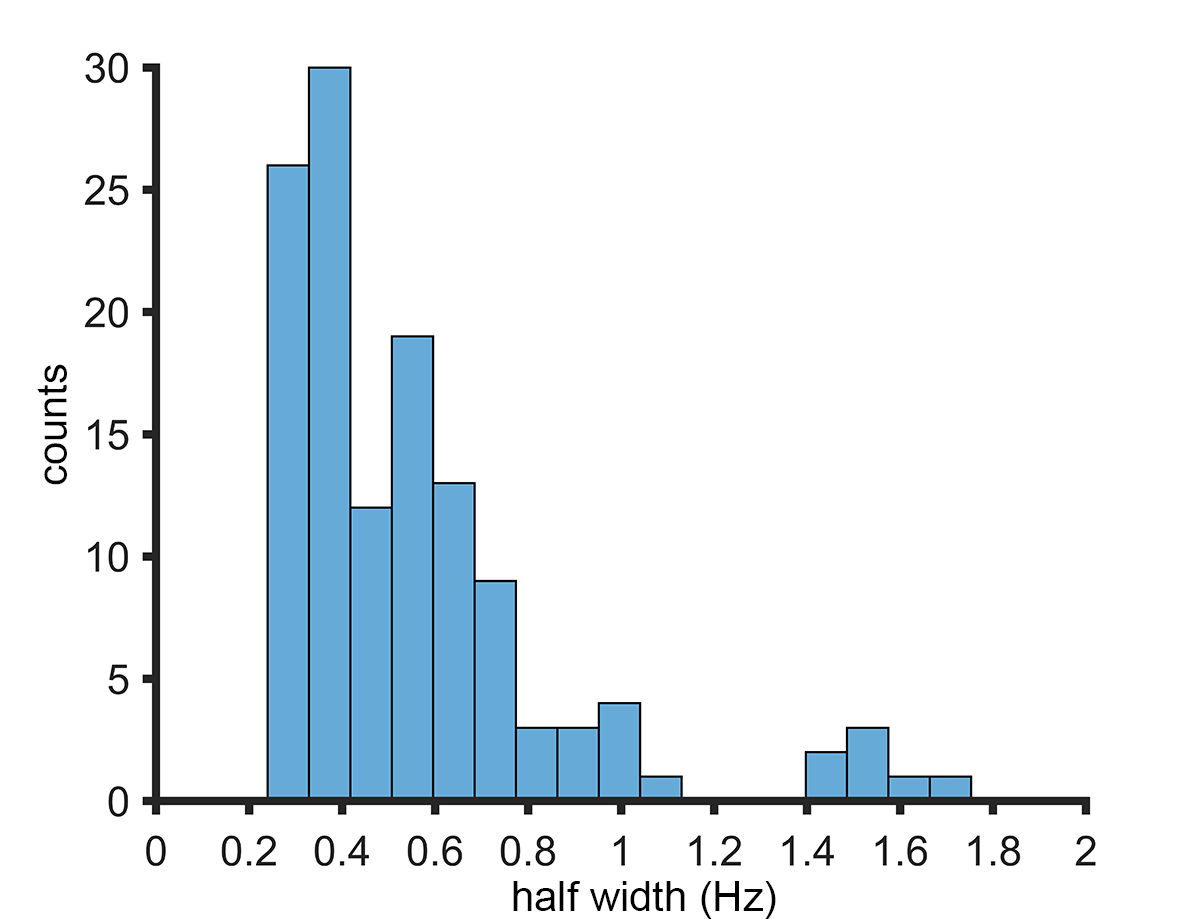

Figure S8: The half width of the peaks indicates a mean half-width of around ~0.5 Hz within 3 s. Resulting from the peak analysis of the simulation of prime number frequency input from 2 - 239 Hz, the half-width of the peaks is demonstrated. The input frequencies were continuously applied. The length of the analyzed signal is 3,000 ms and was applied in the modeled waking state (see **Fig. 2A**). The peak analysis shows that non-local information storage is efficient and works over the whole band-with. Using the equally distributed half-width of the peaks, the coding potential within a distinct time frame can be estimated (~329 bit/s in a bandwidth of 7 - 500 Hz at a single location). The determination of the peaks improves by dividing the frequency space through its own running average (window size = 60 data points).

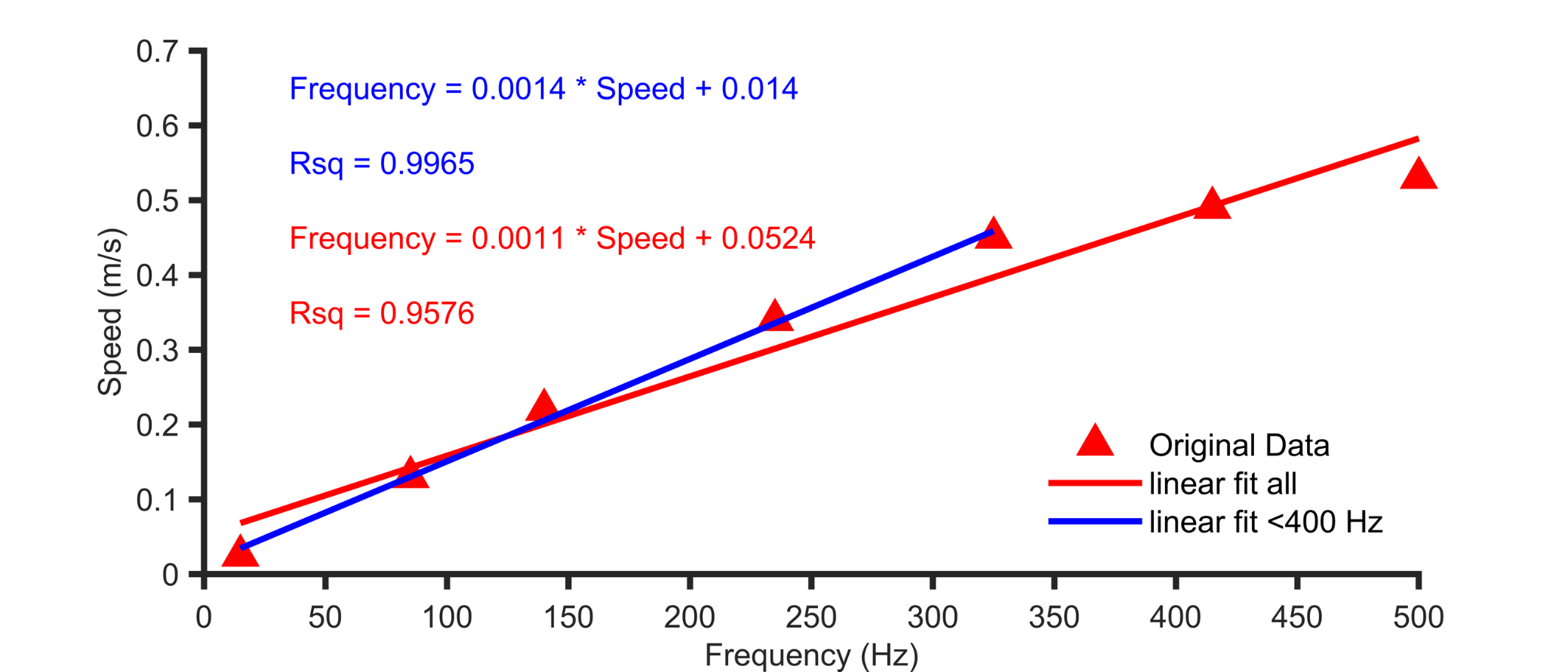

Figure S9: Linear correlation between an increase in wave speed and an increase in frequency. Via the *NI_slopev* energy transfer modulator, the maximal frequency that could be applied to the model was adjusted. The speed at which the wave hills traveled increases linear with increase of the energy transfer (*NI_slopev*) and with the maximal frequency that can be generated. Typically, due to the harmonic generation, the maximal frequency is limited by the highest harmonic. Close to 500 Hz and *NI_slopev* of >2.6, the speed of the simulated waves converges to 0.5 m/s. The wave speed was estimated by 7 Hz continuous peak frequency. Harmonics were generated, this enables the detection of the maximal frequency that the simulation could process. The speed of the wave hills was altered in 7 steps (*NI_slopev* = 0.01, 0.2, 0.5, 1.2, 2.0, 2.5, 2.6655) resulting in wave speeds of 0.03, 0.13, 0.22, 0.34, 0.45, 0.49 and 0.53 m/s. Other parameters were set as follows: *slopeo_damping* of 0.01, *damping* of 0.0001, *marginextradamping* of 2, *inputstrength* of 800 mV and a simulation length of 3 s.

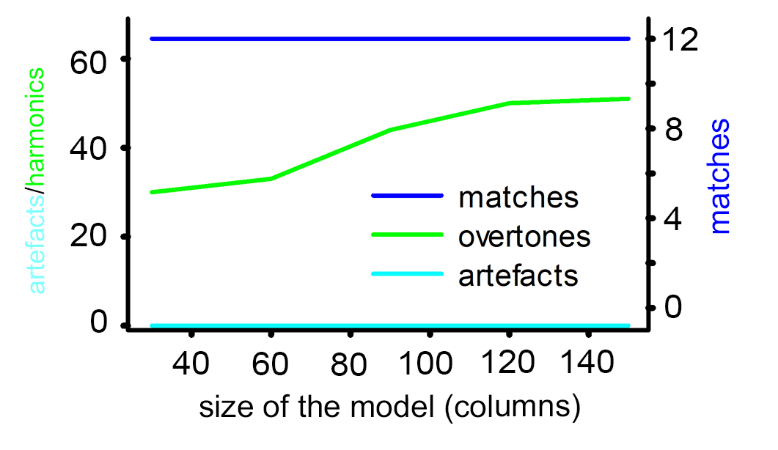

Figure S10: Scalability of model-size - similar wave separation power on direct input but increased self-organization power of harmonics with increased model size. A 20x20 matrix, as well as a 150x150 matrix were equally resolving 12 continuous peak stimuli (29, 31 37, 41, 43, 47, 53, 59, 61, 67, 71, 73 Hz) without generating artificial frequencies. However, increased number of harmonics with model size might be related to increased capability for self-organization, or increased resonance frequencies. The matched frequencies, harmonics, and artificial frequencies were investigated. The x-axis represents the diameter in number of processing units (0.5 mm each). The left y-axis corresponds to the number of detected artificial signals and harmonics. The right y-axis represents the number of input frequencies. Further parameters: *NI_slopev* = 2.6655, *slopeo_damping* = 0.01. *damping* = 0.0001, *marginextradamping* = 2.0 and signal length = 3000 ms. Analysis was executed on MEA signals.

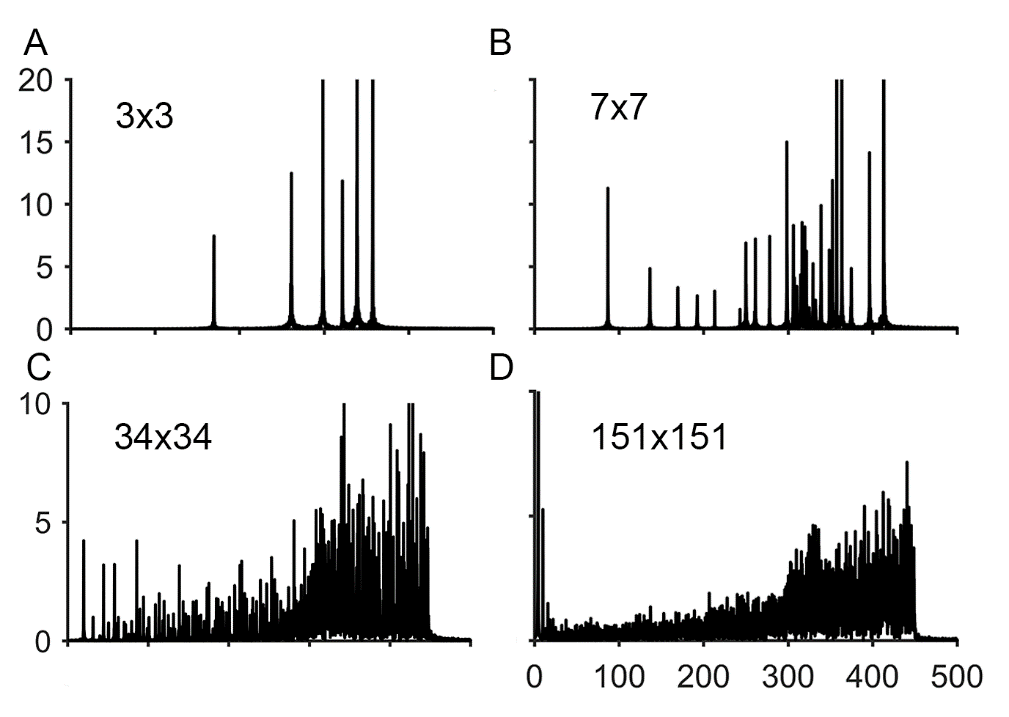

Figure S11: Analyzing the effect of chaotic spontaneous activity on different model sizes shows that the number of resonating frequencies or states increases drastically with model size ((**A**) 3x3, (**B**) 7x7, (**C**) 34x34, (**D**) 151x151 columns). The signal was generated applying spontaneous activity of 100 mV and 0.01 % of activated processing units each time step. Further parameters were as follows: *NI_slopev* = 2.6, *slopeo_damping* = 0.01. *damping* = 0.0001, *marginextradamping* = 2.0 and signal length = 10000 ms. Here, the MEA signals were analyzed.

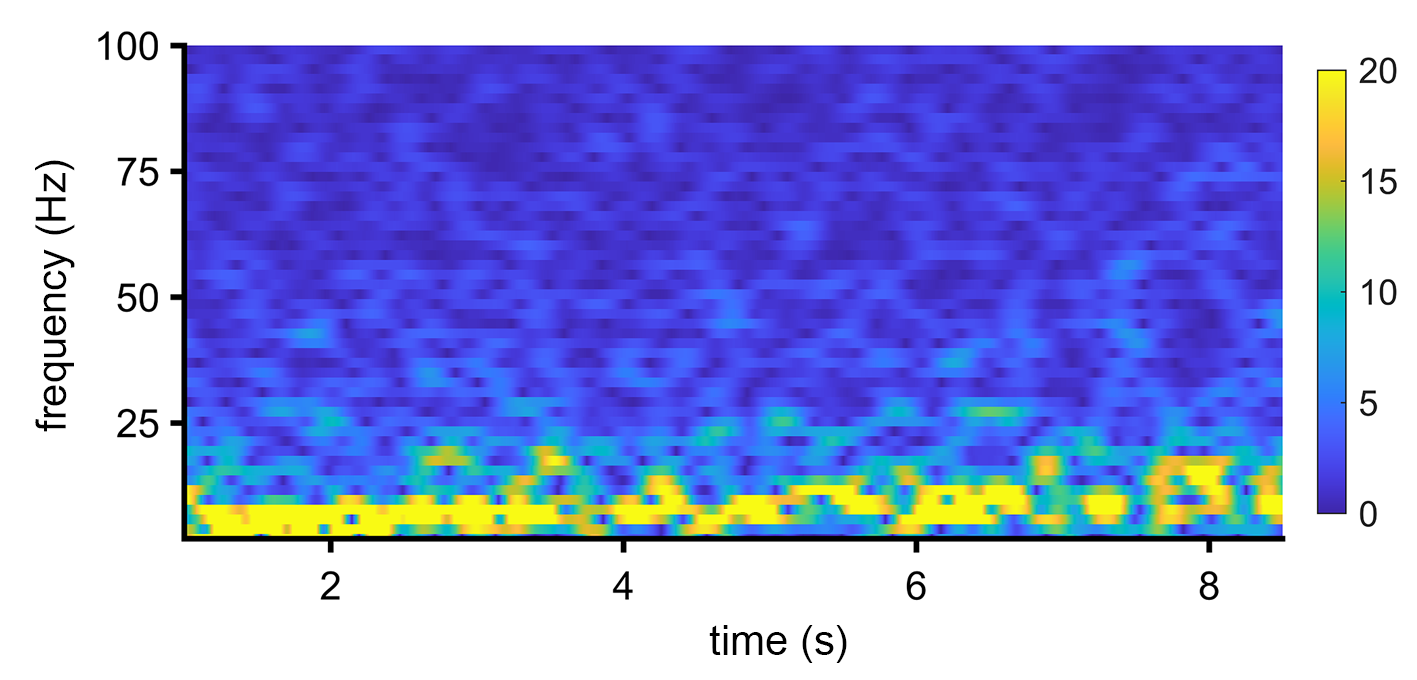

Figure S12: Simulated arousal: Basal brain EEG activity changed from theta (SWS) to alpha activity (waking). For this, the baseline change was mediated in our model by changing the parameter *NI_slopev* (theta to alpha). The *NI_slopev* was stepwise increased from 0.1 to 2.8 by 0.3 each 1 s. The other parameters were the same as in **Fig. 3C**. The signal length was 10 s. Here, the period from 0.5 to 8.5 s is shown. The EEG-electrode radius was 10 mm. The transition from theta to dominating alpha activity in accordance with transition from sleep to waking has been observed in EEG recordings in men ^9^.

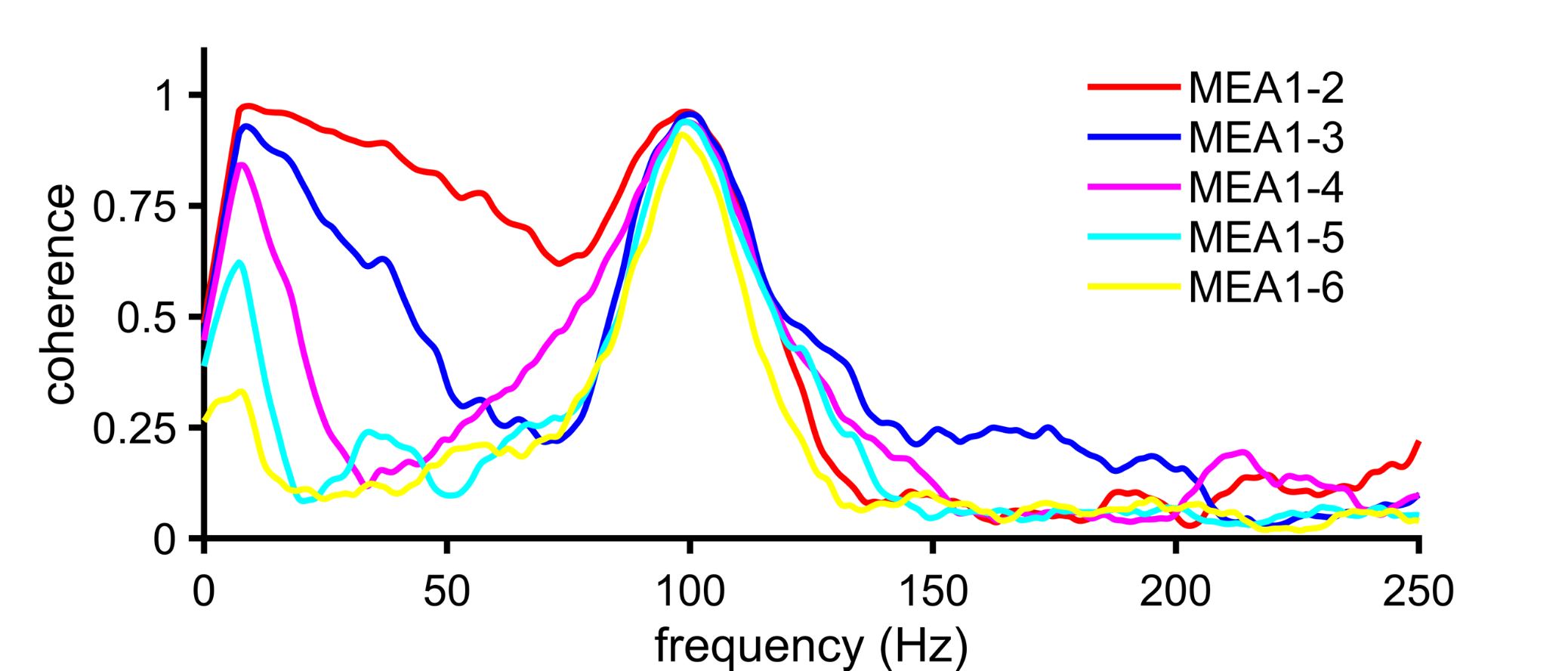

Figure S13: Coherence decreases with space and frequency, but harmonic stimuli can compensate that decline. 500 ms after start of the simulation, a sine stimulus of 100 Hz was computed. The stimulus lasted for 200 ms. The coherence increased at the input frequency. Parameters were the same as in **Fig. 4D**.

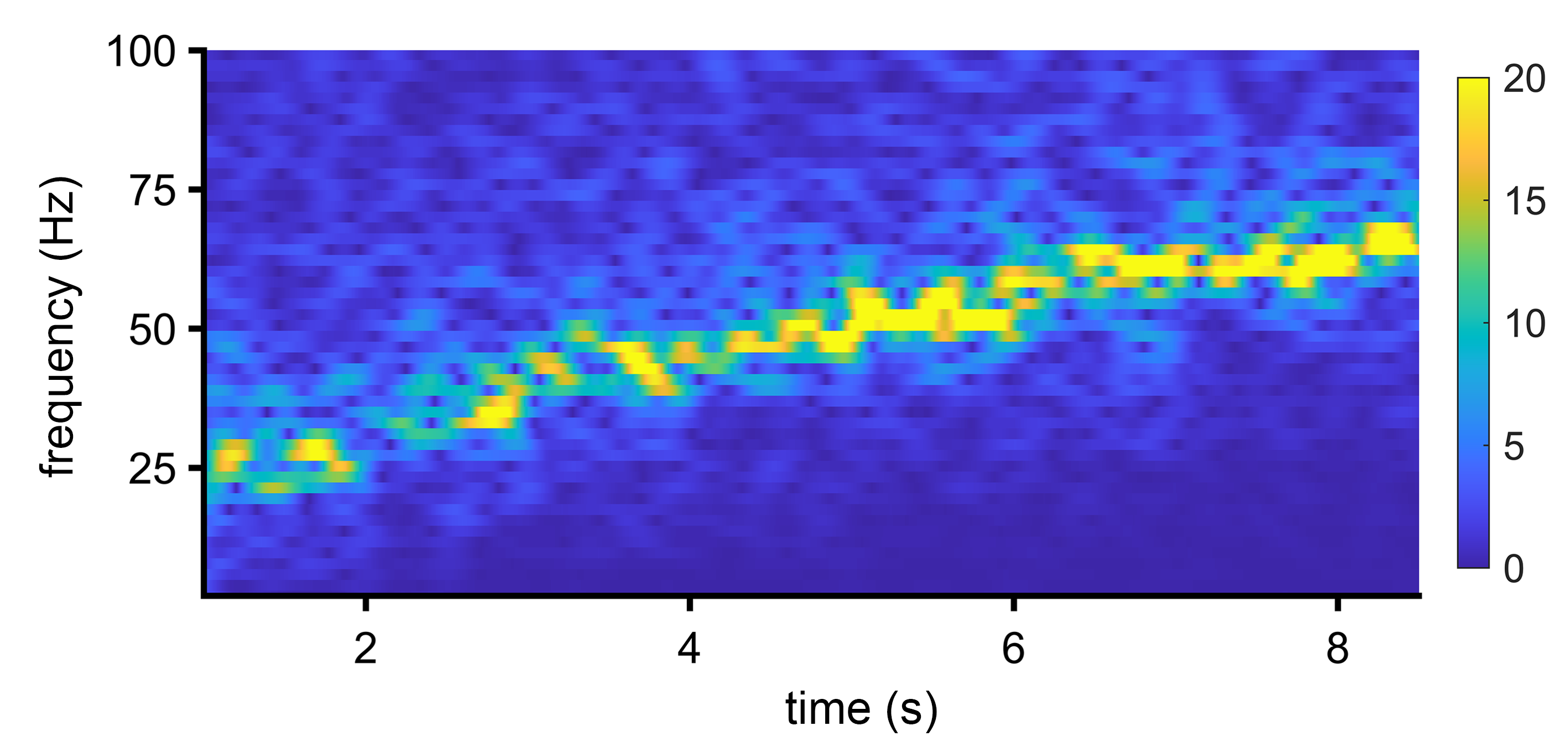

Figure S14: EEG beta firing increases with increased relative activity of inhibitory neurons (increase of *ratio_inhibition_activation1*). Here, we used a similar parameter set up as described in **Fig. 3C**. The *ratio_neighbour_activation* (excitatory neurons) and the *ratio_inhibition_activation1* (inhibitory neurons) were set to 0.8 at time point 0. The *ratio_inhibition_activation1* is increased each second by 0.02. As a result, the self-organized frequency band is increasing gradually. This illustrates that the favored frequency band increases due to more inhibitory neuronal activity in our simulation.

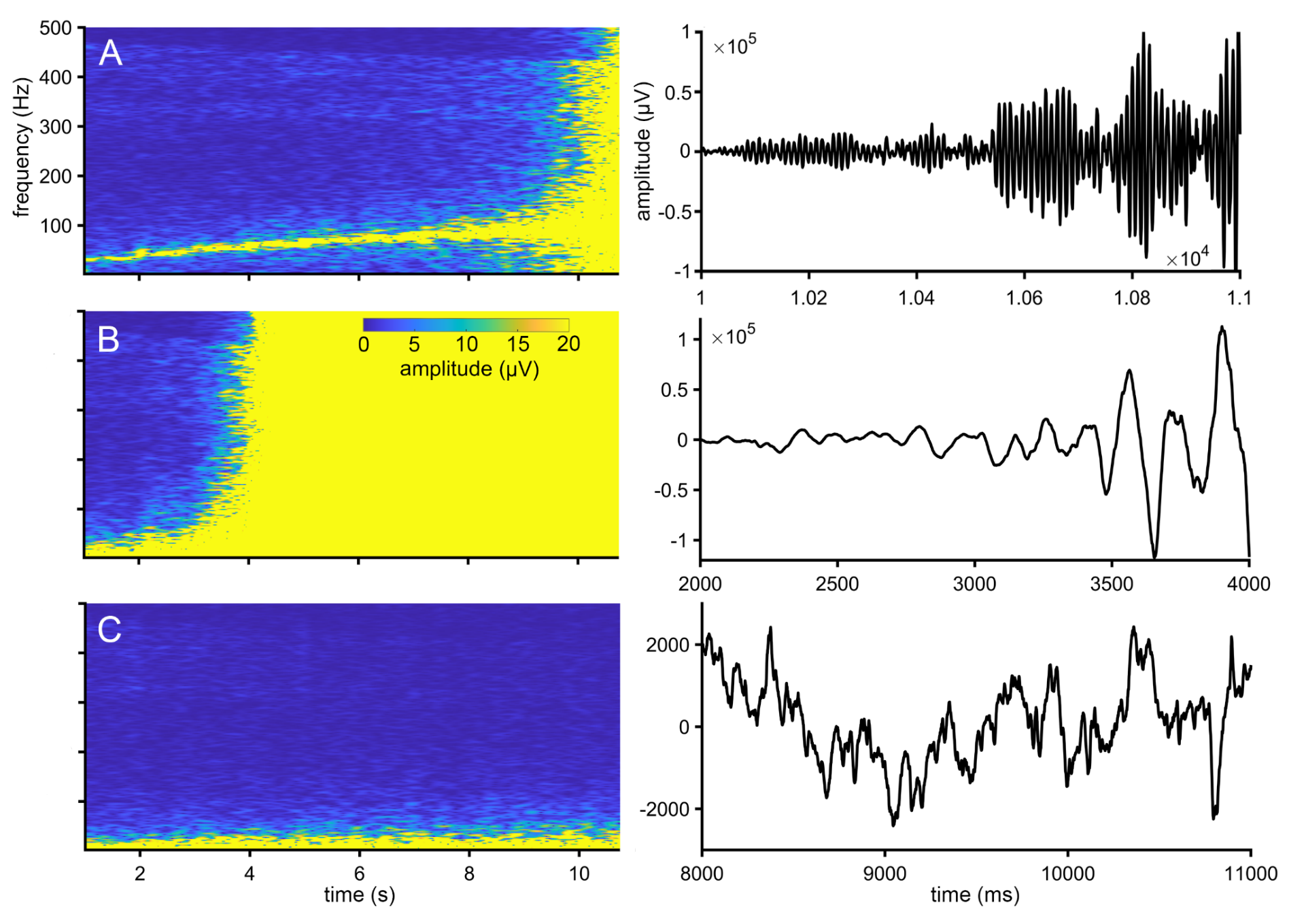

Figure S15: Analyzing epilepsy considering non-local information processing. In **(A)** the *ratio_transmitted_energy* of the neighboring neurons was gradually declined by 0.05 each second and results in a value of 0.5. The process of declining the *ratio_transmitted_energy* was uncorrelated, meaning a random value between, e.g. 1 and 0.5, determines the ratio of energy transfer of the columns. This was analog to an uncorrelated and increased influence of inhibitory neurons. In **(B)** the *slope_vector* and in **(C)** the *slope_old* were statistically declined like in **(A)**. An uncorrelated change of the extent of *slope_vector* and *slope_old* was not disrupting the balance of excitatory and inhibitory neurons, but the balance of integration steps. This could lead to the escalation of signals **(B,C).**

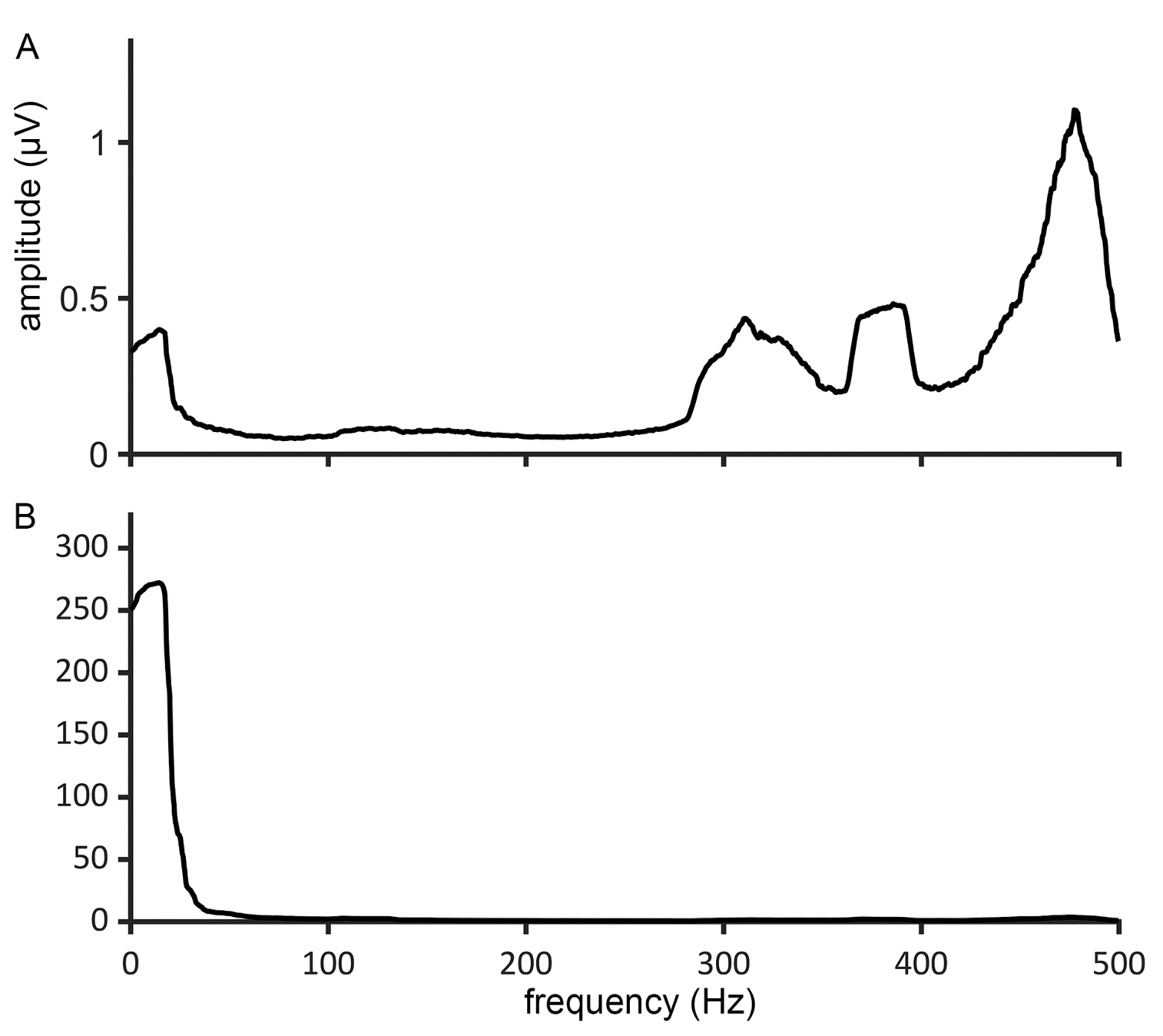

Figure S16: Spatial undersampling suggests that higher frequency bands carry information that is masked by larger electrodes. This effect is illustrated in the following. **(A)** Frequency space of simulated MEA recordings and **(B)** the of simulated EEG recordings. Instead of one signal, 21 different signals per second were randomly applied. The input signals were composed of bursts (>400 Hz) superimposed on carrier waves (<100 Hz). The EEG analysis in frequency space suggests low frequency coding (**B**), whereas MEA points to high frequency coding (**A**). The signal length was 8,000 ms. The signal was smoothed in frequency space with a running average using a window size of 120 data points. Other parameters were the same as describe for **Fig. 2A** for waking.

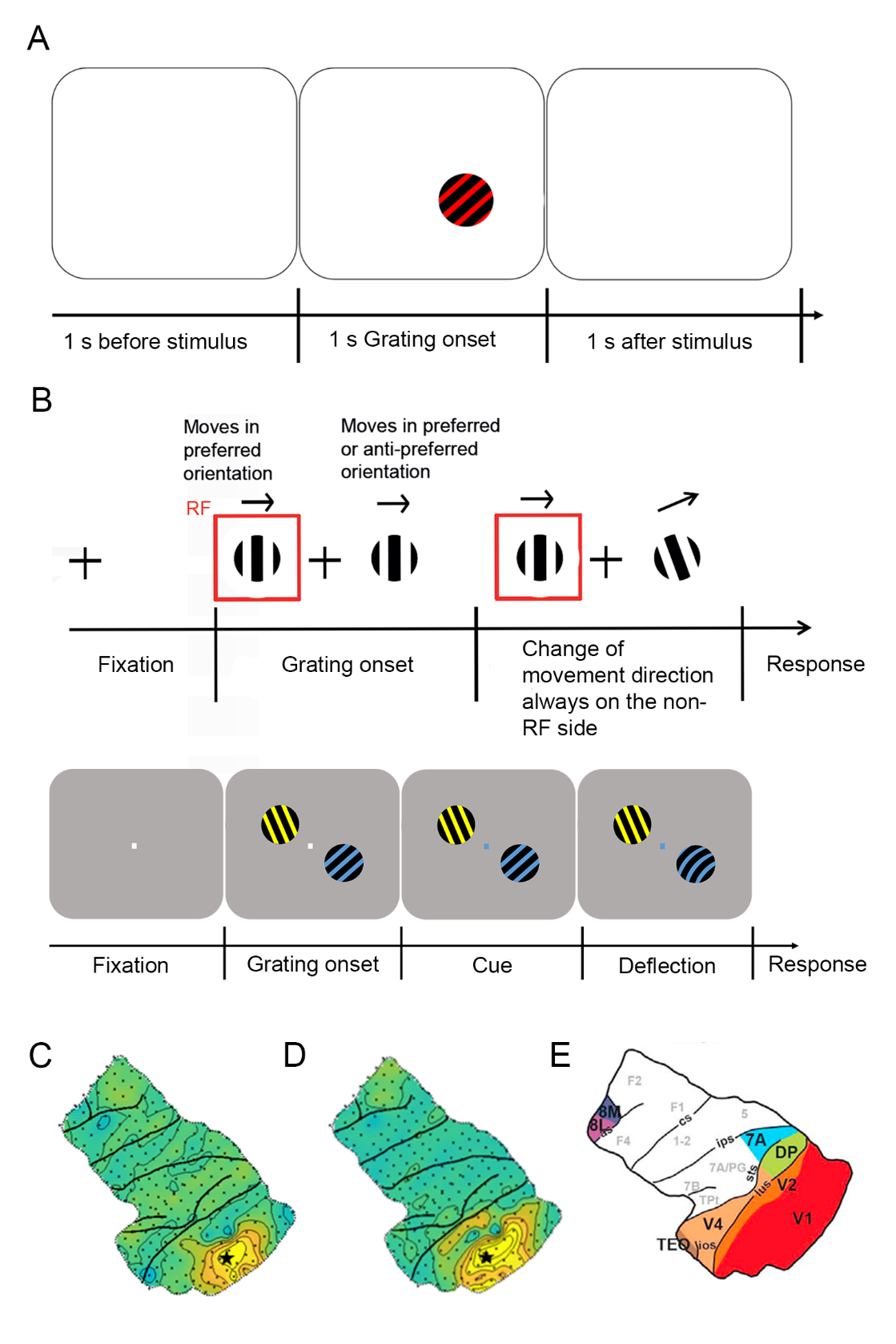

Fig. S17: In silico and in vivo stimulus paradigm. **(A)** The parameters of the grating and the simulated electrodes were chosen to be close to the in vivo experiments. The grating was defined by a harmonic stripe pattern introduced by constant positive (+10 mV, shown in red) and negative (-9 mV, in shown in black) activation levels. The input was permanent, as long as the grating stimuli was on (1 s) and was turned off for 1 s before and after the grating. For the simulation, data was calculated for a small and a big virtual EEG-electrode (radius = 2.5 mm and 10 mm). The small proximate electrode measured direct sensory input at the site of the grating (distance = 0), thus fully overlapping with the stimulus. The larger distant electrode measured at a distance of 15 mm to the stimulus center, thus partly overlapping with the stimulus. The model size was 75x75 mm. Other model parameters were the same as for the simulated waking state (see M&M). **(B)** **Upper paradigm** as used in the V1 microelectrode recordings. A fixation point was presented during a prestimulus baseline (1 s), followed by two luminance gratings (contrast of 100%, diameter of 2–3°, spatial frequency of 1–2 cycles/°, temporal frequency of 1–2°/s). The grating within the receptive field (RF) of the recorded neurons always had the preferred orientation and was moving. The grating outside the RF, on the opposite side of the fixation point, was either presented in the preferred or anti-preferred orientation and was also moving. The monkey had to detect a change of movement direction, which always happened on the side, which was not covered by the RF. **Lower paradigm** used in the ECoG recordings. A prestimulus baseline (0.8 s, fixation) was followed by the presentation of two isoluminant and isoeccentric drifting sinusoidal gratings (diameter: 3°, spatial frequency: ≈1 cycle/deg, drift velocity: ≈1 deg/s, resulting temporal frequency: ≈1 cycle/s, contrast: 100%), one yellow, the other blue. After 0.8–1.3 s, the fixation point changed color indicating which grating was task relevant. At random time points between stimulus onset and 4.5 s after cue onset, either the cued or the uncued grating was slightly bent and the monkey should release the bar on detection of the change. **(C)** The topography shows **induced power (**power values are baseline corrected from -0.25 -0s, relative change) averaged between 200 and 1,000 Hz in the time period of 0 – 0.5 s after stimulus onset) plotted for every electrode of the ECoG grid. The electrode used for **Fig 5** (evoked and induced) is marked by an asterisk **(D)** same as **(C)** but the **evoked power** distribution is shown **(E)** Anatomical landmarks beneath the grid electrodes. The grid was placed over the left hemisphere.

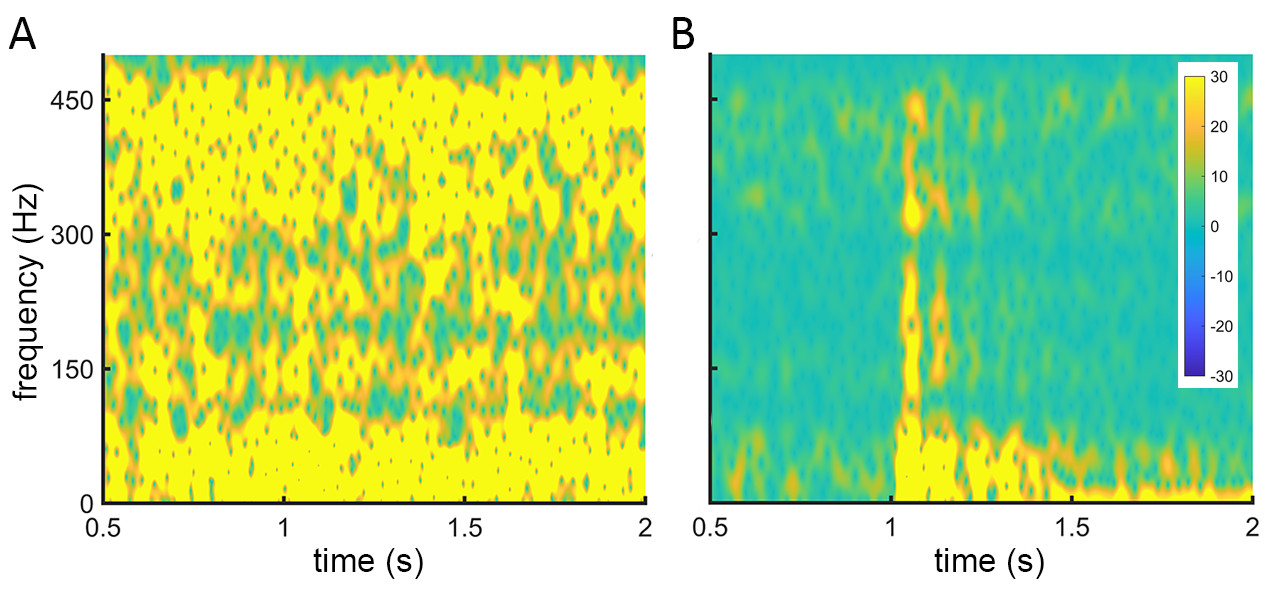

Figure S18: HF and LF coding of a grating signal in the neocolumnar non-local information processing simulation. We demonstrate here the grating signal response with a short-time Fourier transform. The onset of the grating signal was 1 s after trial start. The grating signal lasted for 1 s and a period of 1 s followed after offset of the grating signal (here only 2 s outlined). The trial was repeated 100 times resulting in a signal length of 300 s. 21 short-burst signals per second of random HF and LF coupling, random onset and random frequency were applied to the model to simulate an active background processing model. Decoding the signal of a single trial at a distant stimulus (30 mm from stimulus center) showed a highly loaded coding in frequency space, the induced potential **(A)**. In comparison, 100 trial signals were averaged in time and subsequently transformed to frequency space**.** Stable encoded frequency in time and space were still recognized after the averaging whereas signals with random onset and random location showed interference depletion, the evoked potential **(B)**. It can be noted that constant non-periodic input, such as the grating signal, induced periodic HF and coupled LF coding in the non-local architecture.

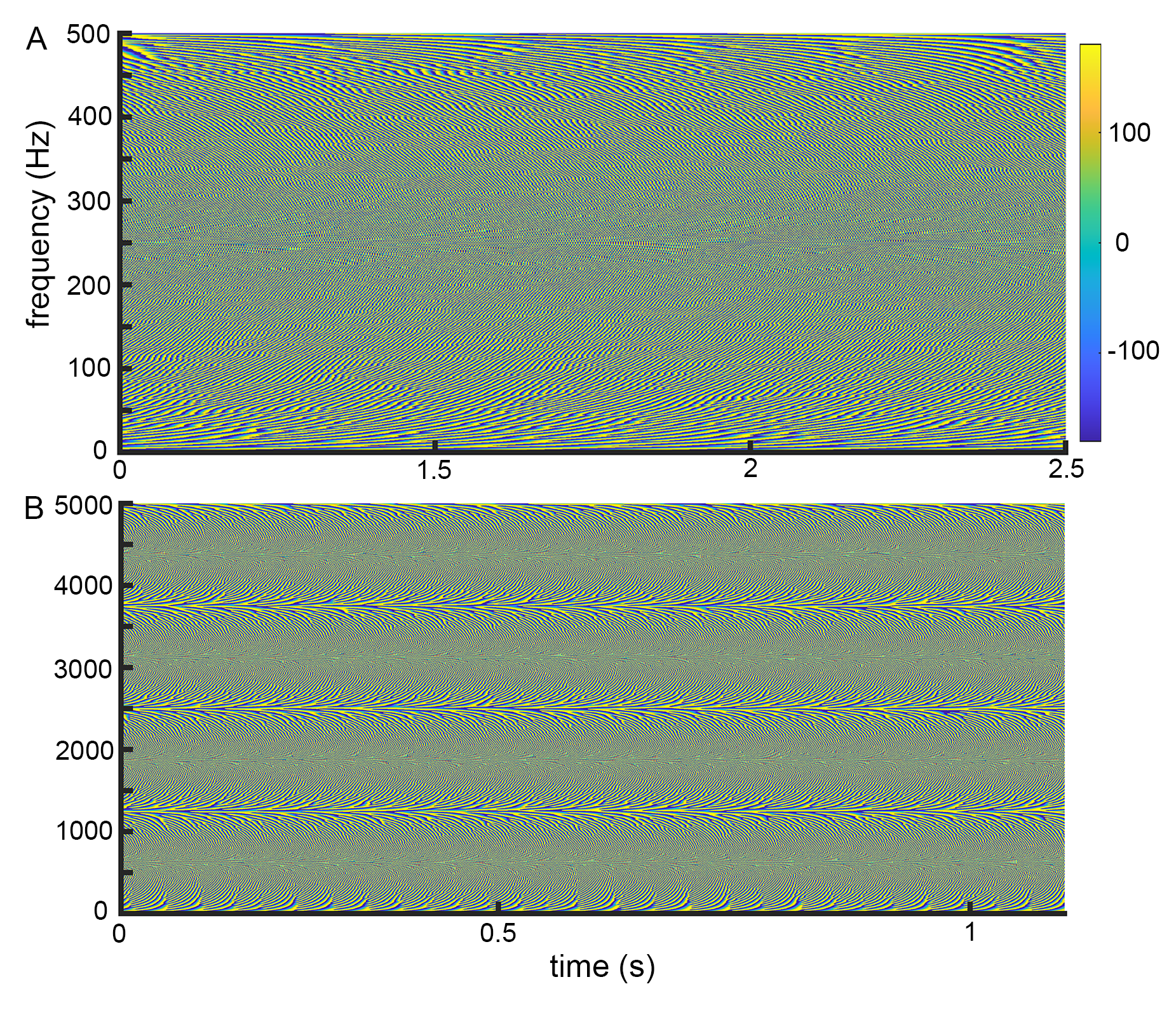

Fig. S19: The phase spaces of signals resulting from the simulation (A) and a recording from a hippocampal brain slice (B) exhibit similar complex patterns suggesting involvement of phase coding. Presented is the angle of the stFT (short-time fourier transform) of **(A)** a baseline signal of the simulation (compare **Fig. 3C**) and **(B)** the baseline signal of *in vitro* field recordings obtained from murine hippocampal slices (compare **Fig. 2D, S6**). The comparable signal pattern of the *in silico* and *in vitro* recordings suggests a homolog processing architecture, following non-local information processing principles.

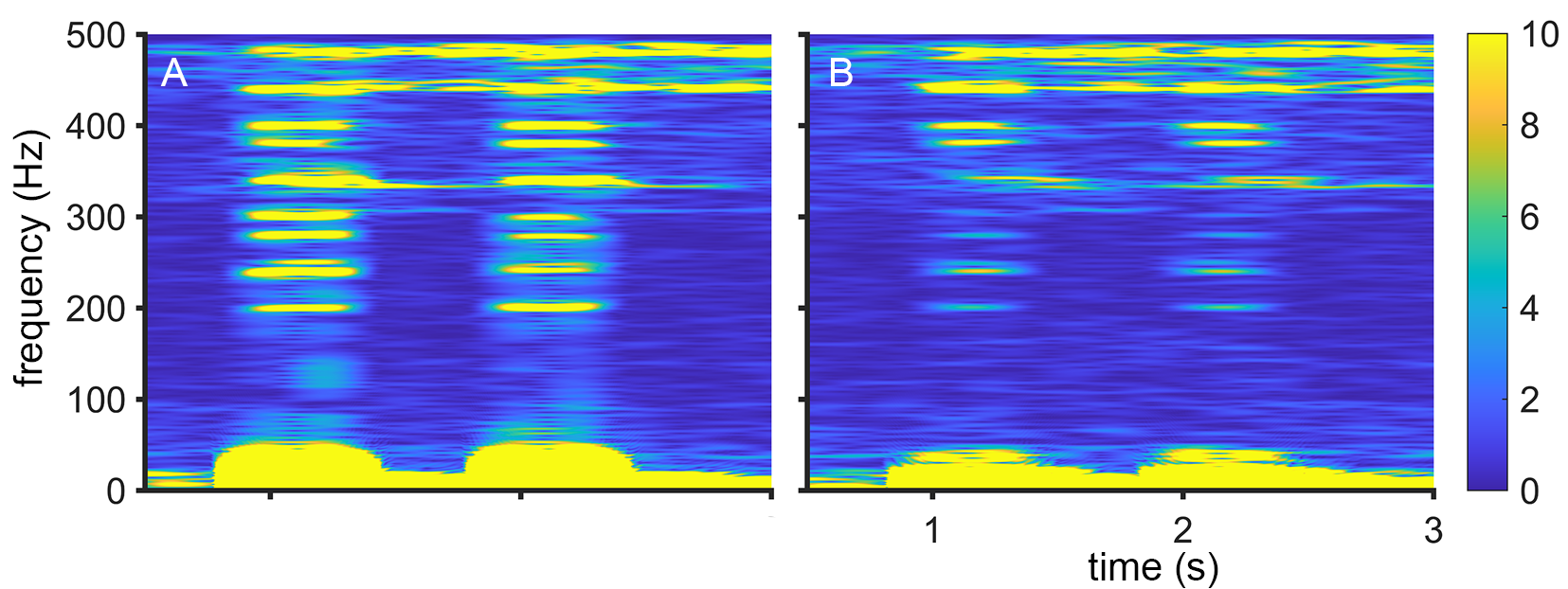

Fig. S20: A phase shift of the certain frequencies could correlate with the distance waves travelled. We analyzed the model processing of an input signal at different distances to the input with a short-time Fourier transform (stFT). Presented is the stFT of EEG1 **(A)** with the coordinates X=2 and Y=2 and EEG3 **(B)** with coordinates X=90 and Y=90. The window size was here 512 ms. The input was applied at location X=1 and Y=1. A Carrier of 10 Hz was applied to 10 bursts of 40-480 Hz (200, 240, 250, 280, 300, 340, 380, 400, 440, 480 Hz) with a burst period of 200 ms length. Burst onset was at 1000 and 2000 ms. The *NI_slopev* was 2.6655, the *damping* was 0.0001, *slopeo_damping* was 0.01 and 0.01 % of the columns were firing spontaneously and chaotic at 50mV. Other parameters were generic. **(A)** All input signals were in phase. There was a continuous activation level <50 Hz, that increased with stimulus onset and carrier wave activity. The frequencies <50 Hz showed the strongest activity, 10 Hz was a true input. Even after a firing period of 1.2 s or 2.2 s theta activity was maintained. **(B)** It is shown that frequencies above 100 Hz have a delayed onset compared to the frequency band <50 Hz and to **(A)**, e.g. the bursts at 240 and 280 Hz are a little bit delayed, the bursts at 300, 380, 400 and 480 Hz are strongly delayed and the burst at 340 Hz is very strong delayed (roughly 300 ms).

###### Table S2: Features of the neocolumnar non-local information processing simulation in context of the literature.

(The table describes simulated and experimental observations and lists content-related references).

| **neocolumnar non-local information processing simulation** | **similarities between simulation and experiment** | **electrophysiological measurements** |
| --- | --- | --- |
| Fig. 1C | Mountcastle ^10^ and Markram et al. ^11^ describe the neocolumnar architecture of the cortex.  Our model architecture is based on the neocolumnar architecture. We simplified columns and interconnected them laterally in order to simulate cortical processing (Fig. 1C). | ^10, 11^ |
| Fig. 1C | Microcircuits of inhibitory and excitatory neurons drive oscillation in cortical regions ^12, 13^.  For simulating the innercolumnar neuronal network of the columns, we simplified the columnar connections to a microcircuit model. In this way we established a microcircuit model of excitatory and inhibitory neurons that are interconnected with each other (Fig. 1). | ^12, 13^ |
| fig. S5 | Chialvo ^14^, Ribeiro *et al.* ^15^ and Fraiman *et al.* ^7^ describe Ising like dynamics in the cortex, such as the criticality. They show that cortical signals show a power law or lognormal distribution which suggests a critical state of information processing.  We could also show the critical distributions in our simulated signals (fig. S5). Furthermore, we intended to compare to the Ising model in more detail and searched for the parameter modulations that enabled chaotic (supercritical) and ordered (subcritical) distributions. The states order and chaos are dominant in the Ising model, but not in the non-local information processing model. In our model the critical distributions dominate (fig. S5). | ^7, 14, 15^. |
| Fig. 1C | Ripples indicate bursts of high frequency activity that are distributed laterally in cortical regions as shown by Oritz et al. ^16^  In the model bursts of high frequency oscillations are a side effect of complex stimuli interference and are distributed laterally (Fig. 1C). Fast ripple-like energy changes are evident in small electrode recordings. | ^16^ |
| Fig. 1C | Evoked signals recorded by big electrodes, e.g. EEG electrodes, often appear like event related potentials (ERP), as shown by Beres ^17^.  In the model, ERP-like signals are simply the sum of signals recorded by small electrodes that are majorly composed of fast ripple-like signals (Fig. 1C). | ^17^ |
| Fig. 2A | We found starting support in literature that neurons can decode certain frequencies ^18-22^, but subsequent research has to be done.  In the model we simply assume frequency coding and decode the signals with a Fourier transform (Fig. 2A). | ^18-22^ |
| Fig. 2B, C and S6 | Ribeiro et al ^15^ analyzed critical distribution of cortical signals in detail. He discriminated distributions of awake, SWS and anesthetized animals.  We looked for critical distributions in our model and could show similar distribution for waking, SWS and anesthesia. Especially, the shift from a lognormal distribution in the waking state to a power law distribution in the anesthesia state could be correlated with the corresponding energy coupling parameter of the simulation (Fig. 2B and C). | ^15^ |
| Fig 2E+S7 | Harmonics appear in cortical signals in response to rectangular input. The harmonics are diminished by sinusoid signals ^23^.  In Fig. 2E we show that harmonics self-organize in response to a periodic peak signals. Further, we could show that sinusoid signals decline harmonics (fig. S7). | ^23^ |
| fig. S9 | A positive correlation of frequency and speed of waves was shown by Zhang and Jacobs ^24^.  An increase of *NI_slopev* increases the maximal frequency that can be processed and both have a positive correlation with the speed of waves (fig. S9). The model wave speeds are comparable to the estimated wavespeeds described by Lubenov and Siapas ^25^. | ^24, 25^. |
| Fig 3C and S12 | A baseline activity of 8 Hz was observed in experiments ^26-28^.  A baseline activity around 8-10 Hz self-organizes in the simulation when spontaneous activity is applied in waking state (Fig. 3C). In the SWS state the baseline activity organizes around <4 Hz (fig. S12). | ^26-28^ |
| Fig. 3D+S13 | The coherence of cortical signals decreases with frequency and distance ^29, 30^.  The coherence of the non-local simulation (Fig. 3D) declines with increasing frequency and interelectrode distance We show that harmonic stimuli can counteract this effect (fig. S13). | ^29, 30^ |
| Fig. 3E | The Lempel Ziv complexity (LZC) was utilized to discriminate between different brain states ^31-34^.  We used the LZC to discriminate between different simulated model states (Fig. 3E). | ^31-34^ |
| Fig 3F | The loss of lesions is well documented ^35-37^ and the robustness to defects is a distinct characteristic of holographic processing ^38^.  AD resolves that the simulation is robust to lesions (defects). The accumulation of defects does not decline coding directly. Even 20 % of defects still allow coding and information is rather localized than fully diminished (Fig. 3F). | ^35-37^ |
| Fig. 3G | Uncorrelated energy transfer can occur on different levels of neuronal processing ^39-42^.  We simulated schizophrenia by introducing uncorrelated energy coupling between the neurons. We observed that the simulation is sensitive to uncorrelated processing and produces artificial frequencies (Fig. 3G). | ^39-42^ |
| Fig. 3H | The evoked gamma-band response is decreased in schizophrenia ^43, 44^.–  In the simulation we show that uncorrelated energy coupling can cause a gamma-band decline (Fig. 3H). | ^43, 44^ |
| fig. S14 | Beta band firing increases with increased GABAergic neuron activity ^13, 45-47^.  In the simulation, the frequency of the self-organized baseline activity increases with increased inhibitory neuron activity (fig. S14). | ^13, 45-47^ |
| fig. S15 | Epileptic seizures may arise when a neuronal network at criticality crosses a sharp border causing activities to escalate ^48, 49^.  In the simulation, we show that overstepping a sharp boundary of maximum energy coupling (*NI_slopev* of ~2.6655) causes energy escalations. Furthermore, we outline that certain modulations of different energy coupling parameters cause different escalating signals (fig. S15). | ^48, 49^ |
| Fig. 4A and 5A | High-frequency coding self-organizes after visual stimulation with a grating. The HF signals is phase-locked with the LF signal. In Fig. 5 we show this response to a grating signal for both, the simulation and *in vivo* experiments in monkey. | Fig. 4B and 5B |

### Additional online information and data visualization

<https://www.biozentrum.uni-wuerzburg.de/bioinfo/computing/neuro>

• PackageNetlogo.rar

- code for our simulation (Netlogo Model)
- tutorial for using the simulation
- Matlab analysis scripts
- example output

• calcium imaging (cell culture experiments) these videos are given directly at the website as well as in the supplementary videos.

• supplementary videos S1-S13, in particular we have

- simulation of anesthesia-like states (Ketamin, Propofol; **suppl. video S8** and **S9**)
- rapid-eye-movement-like sleep state (REM; **suppl. video S10**)
- disease modelling: Alzheimer’s disease (AD; **suppl. video S11** and **S12**) and
   schizophrenia (**suppl. video S13**).
