## Supplementary material for "Modelling non-local neural information processing in the brain": Online Methods

### In silico non-local information processing model: mathematical description

The non-local information processing hypothesis suggests initial uniformity of energy transmission of nodes in a network. The information entering the system is processed and distributed system wide. The processing of the model was discrete in time. The energy change of each time step of each unit was determined by estimating a slope (*slope_vector*) from the difference of the averaged present energy level of the neighbors (*activation1(N1-8)*) and its own present activity (*activation1*). The *slope_vector* was subsequently added to the *slope_old* of the processing unit in focus to update the current state of the slope. The updated state of the slope was added to the *activation1* to generate the current energy level (*activation0*). This represents an integration process with three steps of processing of information.

(step 1) A spatial derivation is calculated with no temporal history.

(step 2) The value of spatial derivation is integrated by adding the derivation to the history of derivation (temporal integration).

(step 3) The updated history of the slope is added to the history of energy levels to determine the current energy state (second temporal integration). The derivation of *activation0* is a temporal derivation and results in a time-dependent function. The second derivation of *activation0* is again a temporal derivation which results in a space-dependent function.

The main algorithm processes information in each node independently and discrete in time. The simulation can be described mathematically as follows:

Assumed are 8 neighbors with activation levels (*activation1(N1-8)*), activation of unit in focus (*activation1/activation0), activation3, activation4*, *slope3, slope4,* *slope_old*, *slope_vector*, and *neighbour_integration* (*NI_slopev*).

#### Main algorithm:

1.	activation3 = (activation1(N1) + activation1(N2) + acttivation1(N3) + activation1(N4))/4

2.	activation4 = (activation1(N5) + activation1(N6) + acttivation1(N7) + activation1(N8))/4

3.	slope3 = activation3 – activation1

4.	slope4 = activation4 – activation1

5.	slope_vector = (slope3 + slope4)/2

6.	slope_old = slope_old + slope_vector

7.	activation0 = activation1 + slope_old

8.	activation_change: within the main algorithm activation1 is the present energy level.

Modulation outside of the main loop is done on activation0. It is important to differ between those equal variables, as the processing units are queried several times within a processing step. The main algorithm results in **equation 1**.

*i* = time steps

*k* = number of neighbors

$$slope\_vector(i)=\frac{\sum_{n=1}^{k} \left( {activation0}_{n}\left( i \right)-activation0(i \right))}{k}$$

$$slope\_old(i+1)=slope\_old\left( i \right)+slope\_vector\left( i \right)$$

$$activation0\left( i+1 \right)=activation0\left( i \right)+slope\_old\left( i+1 \right)$$

$$activation0(i+1)=activation0(i)+slope\_old\left( i \right)+\sum_{n=1}^{k} \left( {activation0}_{n}\left( i \right)-activation0(i \right))/k$$

### *In silico* non-local information processing model: parallel computing

The simulation is written in C, while the benchmarking and analysis script in written in Python. The source-code for both is open-source and available on GitHub: https://github.com/DescartesResearch/BrainSimulation.

We simulated a quadratic grid with increasing node sizes to up to 400,000 nodes in total on four different CPUs. Each measurement was repeated five times per CPU. The different types are a 2-core Skylake laptop (Intel i7-6600U), a 10-core Haswell (E5-2650V3), a 12-core Broadwell (E5-2650V4), and 4-core Skylake architecture (Intel E3-1230V5) system. In total, 970 measurements (194 grid sizes × 5 measurement repetitions) were conducted for each system.

### *In silico* neocolumnar non-local information processing model

For the purpose of setting up a brain slide simulation, Netlogo 6.1.1, a multi-agent programmable environment, was used ^1^. This is a combined approach of programming environmental and visual representation, as well as graphical modulation. Our full custom-written program is available at: <https://www.biozentrum.uni-wuerzburg.de/bioinfo/computing/neuro>

The model architecture is based on a neocolumnar architecture assuming a neocortical column diameter of 500 µm ^2, 3^. The simulation models a cortex area of 60 mm in diameter, corresponding to 14,400 neocortical columns. Due to the hypothesis that neocortical columns consist of units of around 10,000 neurons ^2, 4-6^, the model represents up to 144,000,000 neurons. This is close to the number of columns approximated for the visual cortex of 20,000 columns and the somatosensory cortex of 5,000 columns, with respect to a size of 77 mm and 24 mm in diameter ^7, 8^. The neocortical columns are laterally interconnected. The energies between the units are processed according to the non-local information processing hypothesis. The *slope_vector* represents the transient energy state of synaptic energy transmission, whereas *slope_old* and *activation0/1* rather represent stored energy levels of the cell bodies (all parameters shown in the microcircuit in **Fig. 1A and C**). The information, encoded in frequency and amplitude, is shared between all columns. As every unit is processing the energies and forwarding it to the neighbors in a similar way, the distribution of energies appears wave-like. In consequence, information can be read out for every unit of the model.

The wave-like distribution of the information can be used to simulate signals that are comparable to signals measured by microelectrodes or EEG-electrodes. The output of simulated signals is given by a spatial and temporal integration of several columns, representative for the size or integrative capacity of a virtual electrode.

The smallest unit measured by a simulated microelectrode is one neocortical column. Therefore, a simulated microelectrode simulates the energy changes within a diameter of 500 µm. When we arrange six of such microelectrodes in a grid, we define this microelectrode grid as MEA (analog to a multi electrode array). The interelectrode distance of the MEA can be adjusted (default setting: 10 mm distance between electrodes).

To mimic LFP-like signals, the activities of neurons within a radius of 1 mm are summed up. Therefore, the diameter of the integrated field of a simulated microelectrode, measuring a simulated LFP signal, is set to a *in silico* value of 2 mm ^9-11^. Signals simulated by an EEG-like electrode are set to an approximated number of around 1,300 columns resulting in an EEG-electrode radius of around 10 mm. The values are a rough estimate and are in line with earlier considerations ^12, 13^. In the program, signals measured by virtual electrodes are displayed in 12 plots (6 MEA-plots and 6 EEG-plots) (see <https://www.biozentrum.uni-wuerzburg.de/bioinfo/computing/neuro>). In the output plots, the time is plotted against the activation (energy) level (in µV). The simulated signals are analyzed by a Fourier transform. The bandwidth of the signals can be analyzed in the range of 0–500 Hz.

Analyzes throughout the manuscript were computed with Matlab R2021a (MathWorks, <http://www.mathworks.de/products/matlab/>). The Matlab analysis scripts are again provided at: <https://www.biozentrum.uni-wuerzburg.de/bioinfo/computing/neuro>.

### *In silico* neocolumnar non-local information processing model: modulation

The modulation of the energy coupling of the components of **eq. 1** can be used to simulate different processing states (e.g. waking states and disease states).

The following steps show the modulation of steps 1-8 of **eq. 1** in method section:

9.	slope3 = (activation3 * ratio_neighbour_activation – activation1 * ratio_inhibition_activation1) * neighbourintegration

10.	slope4 = (activation4 * ratio_neighbour_activation – activation1 * ratio_inhibition_activation1) * neighbourintegration1

11.	slope_vector = (slope3 + slope4) / nc

12.	slope_vector = slope_vector * NI_slopev

13.	slope_old = slope_old / (1 + slopeo_damping)

14.	activation0 = activation / (1 + damping)

15.	activation0 = activation0 / (1 + damping + (marginextradamping / (1+ minimum_margin)))

*Marginextradamping* increases with distance of margin to the center of the model.

The resulting modulated main algorithm (modulation, steps 9-15) is given in **eq. S2**.

**Equation S2**

$$slope\_vector(i)=\frac{\sum_{n=1}^{k} \left( {activation0}_{n}\left( i \right)*{ratio\_neighbour\_activation}_{n}\left( i \right)-activation0(i \right)*ratio\_inhibition\_activation1\left( i \right))}{k}*NI\_slopev$$

$$slope\_old(i+1)=(slope\_old(i)+slope\_vector(i))/(1+slopeo\_damping)$$

$$activation0\left( i+1 \right)=(activation0\left( i \right)+slope\_old(i+1))/(1+damping)$$

$$activation0\left( i+1 \right)=(activation0\left( i \right)+(slope\_old(i)+\sum_{n=1}^{k} \frac{{activation0}_{n}\left( i \right)*{ratio\_neighbour\_activation}_{n}\left( i \right)-activation0\left( i \right)*ratio\_inhibition\_activation1\left( i \right)}{k}*NI\_slopev)/(1+slopeo\_damping))/(1+damping)$$

**Explanation of the code lines**

In the model, the influence of neighbors on the *slope_vector* in the simulation is typically 1/8 (simplification: direct neighbors on a grid). This symmetry can be changed by the influence of neighbor energy (*ratio_neighbour_activation*) and the subtraction of the own energy (*ratio_inhibition_activation1*). The contribution of the calculated *slope_vector* on the update of *slope_old* is modulated by *NI_slopev*. This means that the activity of 8 neighbors and its derivation from the own activity can be summed and transmitted to the central neurons or normalized by the extent of *NI_slopev*.

Likewise, the contribution of *slope_old* to *activation0* can be modulated and the final energy levels are regulated by *damping*.

We assume here a symmetrical calculation of the neighbor energies. However, asynchronous, and individual modulation is possible in the simulation. The translation of the modulation into a microcircuit model is shown in **Fig. 1C**. The modulation on the core steps of non-local processing might function as modulation of the cell body energy state (*damping* and *slopeo_damping*), or the modulation of synaptic transmission (*NI_slopev*).

The modulation by *NI_slopev* is important for the spatial derivation in step 9 and 10 of **eq. 1**, as it represents a symmetrical substraction of *activation1* from the neighboring *activation1* (here named as *activation3* or *activation4*). Altering this symmetry by *ratio_neighbour_activation*, or *ratio_inhibition_activation1* results in resonance catastrophy , as indicated in **fig. S15.**

To mimic different brain states, the values for *NI_slopev, slopeo_damping* and *damping* are modulated. In case of asymmetric modulation of energies by *NI_slopev, slopeo_damping* and/or *damping* systemic, diseases such as schizophrenia, AD, or epilepsy can be simulated.

The schizophrenia simulation was implemented by introducing uncorrelated synaptic information transfer. The transmission of the *slope_vector*, *slope_old,* and/or *activation0/1* was decorrelated randomly.

The AD model was set up by generating randomly deleting information transfer of entire functional units (lesions). The lesion size can be altered by the user and thereby impair information distribution over the entire model.

Epilepsy is simulated by provoking resonance collapse of the model. This can be caused by changing the energy coupling parameters *NI_slopev, slopeo_damping, damping, ratio_neighbour_activation,* and/or *ratio_inhibition_activation1*, or by increasing uncorrelated information transfer.

### Hippocampal brain slice recording

Data from ^14^ are analyzed comparing carbachol induced gamma oscillations in wt mice and a constitutively active form of the erythropoietin receptor in GABAergic neurons **(VEPOR^+/+^)** ^14^. Field oscillations were induced with 20 μM carbachol (Sigma–Aldrich) in transverse 400 μm hippocampal slices from male mice (27–33 day‐old), protocols in ^15^. Extracellular recordings were performed in the *stratum radiatum* of CA3 at 33°C in a Haas‐Top interface chamber, using a MultiClamp 700B amplifier. Data were sampled at 10 kHz using an Axon Instruments Digitizer 1440A with pClamp 10 software (Molecular Devices, Sunnyvale, CA, USA). The analysis was equivalent to **Fig. 2B and C** for baseline activity prior to carbachol application and for 0-15, 15-30, 30-45 and 45-60 min time segments after carbachol application. Results are displayed in **Fig. 2D and S6A (wt)** and **S6B (VEPOR^+/+^)**.

### Grating Experiments

The *in vivo* grating experiments are compared to the *in silico* data generated from the model.

### *In silico* grating stimulation of the neocolumnar non-local model

The grating simulation incorporated a 3 s time period presented 100 times. The grating was defined as continuous input over a stimulation period of 1 s. Before grating onset and after grating offset a 1 s period without input was added. The grating had a radius of 10 mm and had four stripes with a constant input of 10 mV and four stripes with a constant input of -9 mV (compare **fig. 17A** and **suppl. video S14**). As this was a first attempt to simulate experimental input as used in *in vivo* experiments and test basic response properties of our model, we chose a simple non-moving grating. The simulations were done for two types of virtual EEG-electrodes. The center of the simulated proximate EEG-electrode, with a radius of 2.5 mm, was at the same location as the center of the grating stimulus. The center of the distant electrode, with a radius of 10 mm, was located 15 mm apart from the center of the grating stimulus, thus overlapping by 5 mm with the stimulus. The size of the model was 75 x 75 mm (150 x 150 columns) and the energy coupling parameters were set to *NI_slopev* of 2.6655, *damping* of 0.0001, and *slopeo_damping* of 0.01. The full length of a grating experiment was 300 s, including 100 grating onsets (trials).

The analysis followed as closely as possible the analysis of the *in vivo* data, which is described in detail below. There are two exceptions: one, as the sampling frequency of the data was 1000 Hz, the range of frequency analysis was limited to 500 Hz. The high frequency power therefore was assessed for target frequencies between 100 to 500 Hz for shifting time windows over 100 ms in steps of 10 ms. Two, since there was no delay between stimulus onset and model response, we chose the time window for baseline correction to be between -0.25 to -0.1 s.

### *In vivo* grating experiment: V1 microelectrode recording in macaque monkey

All procedures were approved by the ethics committee of the Radboud University, Nijmegen, NL.

Behavioral paradigm and visual stimulation. A trained male macaque monkey participated in a study using a change detection paradigm. Trials started when the monkey touched a bar to start the experiment. A fixation point (~0.2° diameter) lit up and gaze had to be held within a small window around the fixation point, like previously described ^16^. Consecutively, a pre-stimulus baseline (1 s) started, followed by the appearance of two stimuli placed in different visual quadrants (**Fig. S17**). We refer to this time point as stimulus onset. The stimuli were luminance gratings with a contrast of 100%, a diameter of 2–3°, a spatial frequency of 1–2 cycles/°, and a temporal frequency of 1–2°/s. The grating within the receptive field (RF) of the recorded neurons always had the preferred orientation and was moving. The grating outside the RF, on the opposite side of the fixation point, was either presented in the preferred or anti-preferred orientation and was also moving. The monkey had to detect a change of movement direction, which always happened on the side, which was not covered by the RF. Stimuli were presented on a 120 Hz CRT monitor. For the presented analysis, all trials were pooled independent of correct or incorrect responses and preferred or anti-preferred orientation outside the RF. The receptive fields were mapped in separate sessions, beforehand.

Surgery and recording. Electrophysiological recordings were obtained from six to eight tungsten electrodes positioned in V1. Each electrode’s signal was passed through a headstage (Plexon) amplified by a factor of 20. Signal acquisition, filtering, and amplification were done with a Neuralynx Digital Lynx acquisition system. This signal (sampling rate: 32,556 Hz) was used for further analysis.

Analysis. The analysis was done using MATLAB (MathWorks) and the fieldtrip toolbox ^17^.

Preprocessing. Line noise and the harmonics were removed using the discrete Fourier transform (dft) for every analysis except spike detection. Data was padded to 5 seconds for dft and later cut into trials +/- 1.5 seconds around stimulus onset. The trial was demeaned, and artifacts were removed by visually inspecting the variance over trial and time. Whole trials were removed. We afterwards analyzed the 20 sessions with a total of 4,863 trials (mean of 243.15 trials per session, SD: 103.52). The data was further analyzed in 6 different ways:

Local field potential (LFP): To get the LFP, data was lowpass filtered (Butterworth IIR filter, zero-phase forward and reverse filter, filter order 6) at 500 Hz.

Multi-unit activity (MUA): For MUA, we followed an approach similar to the one described in ^18^ and bandpass filtered (Butterworth filter, zero-phase forward and reverse filter, filter order 6) the data between 750 and 8,000 Hz. For plotting, we calculated the absolute values of the Hilbert transform of the filtered data.

High frequency power (induced): The high frequency power was assessed using a FFT based time-frequency analysis in combination with Slepian tapers (discrete prolate spheroidal sequences) to spectrally smooth over a frequency range of +/- 50 Hz around each target frequency. Frequencies targeted were 100 to 1,000 in steps of 100 Hz. The frequency analysis was done in shifting time windows (namely over 50 ms in steps of 1 ms). Time frequency representation (TFR, **Fig 4A**) is plotted from 200-1,000 Hz and baseline corrected (relative baseline change, baseline from -0.25 -0 s) plotted from -0.25 to 0.5 s in steps of 0.025 s. The plotted time refers to the center of the time window.

High-frequency power of the time-domain averaged data (evoked): The same approach was applied for the high frequency assessment of the averaged data, however here, trials were first averaged in the time domain (i.e. mean over the trials within one session), and then frequency demodulated in a time resolved fashion (all parameters as described for the non-averaged data). Time frequency representation (TFR, averaged over all sessions, **Fig 4B**) is plotted from 200-1000 Hz and baseline corrected (relative baseline change, baseline from -0.25 -0 s) plotted from -0.25 to 0.5 s in steps of 0.025s. The plotted time refers to the center of the time window.

Gamma power: Gamma power was assessed using the same time-frequency analysis, however, now using a single Hanning taper with the frequency of interest centered at 60 Hz.

Spiking activity: Spikes were detected with a simple threshold approach. They were first highpass filtered at 500 Hz (Butterworth filter, zero-phase forward and reverse filter, filter order 6), and a positive and negative threshold was determined for each recording session separately. A spike was detected at the maximum absolute point crossing the threshold and cut out +/- 2 ms and plotted for visually checking the detection quality. The data was then converted to binary data (with a one at the absolute maximum of the detected spike) and summed over the same time period as used for the other approaches (namely over 50 ms in steps of 1 ms).

Plot of time series: Time resolved data (i.e. spiking, MUA, LFP, gamma power, 500 Hz power, averaged 500 Hz power) is plotted from -0.2 s to +0.5 s around stimulus onset for a single session (**Fig 4C**). Data was normalized by (mean-min)/(max-min) to be able to show the temporal relationship between the different measures.

Phase-amplitude coupling: The time resolved normalized power in the 500 Hz band (as plotted in **Fig 4C**; 500 Hz power and averaged 500 Hz power) was frequency demodulated by means of FFT (Hanning window, zero-padded to 3 seconds). For **Fig 4D**, the plotted standard error results from the different experimental sessions.

### *In vivo* grating experiment: ECoG recording in macaque monkey

All procedures were approved by the ethics committee of the Radboud University, Nijmegen, NL.

Visual Stimulation and Attention Paradigm. Two trained male macaque monkeys participated in a change detection task. Please note that these monkeys were different from the one participating in the V1 recording and that only data from one monkey (P) was analysed. The experiment consisted of several conditions and a detailed description of the full experiment can be obtained from ^19, 20^ as well as ^21^). As for our particular analysis only one condition was used, the description will focus on this condition (**Fig. S17**). Additionally please note that all trials were pooled independent of the behavioral outcome. Each trial started when the monkey touched a bar while a gray fixation point of 1 degree was shown at the center of the screen. This prestimulus baseline lasted 0.8 s. Then, two isoluminant and isoeccentric drifting sinusoidal gratings were presented, one in each visual hemifield (diameter: 3°, spatial frequency: ≈1 cycle/deg, drift velocity: ≈1 deg/s, resulting temporal frequency: ≈1 cycle/s, contrast: 100%). This is what we refer to as stimulus onset. In any given trial, one grating was tinted yellow, the other blue. These equiluminant colors were randomly assigned across trials. After 0.8–1.3 s, the fixation point changed color. This color served to indicate which grating was task relevant. At random time points between stimulus onset and 4.5 s after cue onset, either the cued or the uncued grating was changed and the monkey should release the bar on detection of the change. The stimulus change consisted of a gentle bend of the stripes. Stimuli were presented on a 120 Hz CRT monitor (same as described above).

Surgery and recording. Neuronal recordings were made from the left hemispheres through a micromachined 252-channel electrocorticogram-electrode (ECoG) array implanted subdurally ^22^. The ECoG was placed directly onto the brain under anesthesia. For details on the procedure, please refer to ^19^. Signals were obtained from the 252 electrode grid and were amplified 20 times by eight Plexon headstage amplifiers, then low-pass filtered at 8 kHz and digitized at 32 kHz by a Neuralynx Digital Lynx system.

Analysis. The analysis was done using MATLAB (MathWorks) and the fieldtrip toolbox ^17^.

Preprocessing: Line noise and the harmonics were removed using the discrete Fourier transform in all analysis except spike number. Data was padded to 5 seconds for dft. Data was cut into trials 0.25 seconds before and 2.15 seconds after stimulus onset. Trials were demeaned. Artifacts were removed by visually inspecting the variance over trial and time. Whole trials were removed (37 out of 104; keeping 73 trials for further analysis).

High frequency power (HF). The high frequency power was assessed using a FFT based time-frequency analysis in combination with Slepian tapers (discrete prolate spheroidal sequences) to spectrally smooth over a frequency range of +/- 50 Hz around each target frequency. Target frequencies were 100 to 2000 in steps of 100 Hz (model: 100 : 500 Hz in steps of 10). Data was padded to 3 seconds. The frequency analysis was done in shifting time windows (namely over 50 ms in steps of 1 ms, model: 100 ms in steps of 10 ms). Time frequency representation (TFR, **Fig 5A**) is plotted from 200-1000 Hz (200-500 Hz for the model) and baseline corrected (relative baseline change from -0.25 -0 s, model: -0.025- -0.1 s) plotted from -0.25 to 0.5 s in steps of 0.025s (model, insteps of 0.01s). The plotted time refers to the center of the time window.

High frequency power of the time-domain averaged data The same approach was applied for the high frequency assessment of the averaged data, however here, trials were first averaged in the time domain, and then frequency demodulated in a time resolved fashion (all parameters as described for the non-averaged data). Time frequency representation (TFR, **Fig 5B**) is plotted from 200-1000 HZ and baseline corrected (relative baseline change from -0.25 -0 s) plotted from -0.25 to 0.5 s in steps of 0.025s. The plotted time refers to the center of the time window.

Topography: For the topography (**Fig S17C and D**), we plotted the power values (baseline corrected from -0.25 -0s, relative change) averaged between 200 and 1000 Hz in the time period of 0 – 0.5 s after stimulus onset). The anatomical location of the electrodes is shown in **Fig S17E**.

Plot of time series: Time resolved data (i.e. 500Hz power, averaged 500 Hz power; both +/- 50 Hz due to the use of Slepian tapers as described above) is plotted from -0.2 s to +0.5 s around stimulus onset for a single session (**Fig 5C**). Data was normalized by (mean-min)/(max-min) to be able to show the temporal relationship between the different measures. The 1000 Hz sampling frequency resulted from the 1 ms shift of 50 s analysis windows as described above for the power analysis.

Phase-Amplitude coupling. The time resolved normalized power in the 500 Hz band (as plotted in **Fig 5B**; 500Hz power and averaged 500 Hz power) taken from -0.2 to 0.5 sec after stimulus onset was frequency demodulated by means of FFT (Hanning window, zero-padded to 3 seconds). For **Fig 5D**, power values were averaged over all trials; only the non-averaged power values have an SE, which comes from the trials.

### Calcium imaging and analysis

The neuronal calcium activity **videos S1** and **S2** were processed using the Neural Activity^3^ (NA^3^) software by Prada et. al ^23^. This software detects calcium activity peaks based on a continuous wavelet transform (wavelet ridge walking). First, the image field of the video is divided in a grid, each square in the grid is analyzed as an independent calcium trace. The wavelet transform is applied to each of the traces, using the mexican-hat as mother wavelet. From the spectrum of each trace, the ridges are located and organized in a tree structure according to their vicinity in the trace. Later those ridges are filtered leaving out the ones corresponding to noise and keeping only the ones that contain possible activity peaks. Finally, the highest peak on each of the ridges is accounted as an activity peak according to its signal-to-noise ratio.
